# Conditional Requirement for Dimerization of the Membrane-Binding Module of BTK

**DOI:** 10.1101/2023.10.05.561114

**Authors:** Timothy J. Eisen, Sam Ghaffari-Kashani, Jay T. Groves, Arthur Weiss, John Kuriyan

## Abstract

Bruton’s tyrosine kinase (BTK) is a major drug target in immune cells. The membrane-binding pleckstrin-homology and tec-homology (PH–TH) domains of BTK are required for signaling. In vitro, dimerization of the PH–TH module strongly stimulates BTK kinase activity. Whether BTK dimerizes in cells via the PH–TH module, and whether this dimerization is necessary for signaling, is unknown. To address this question, we developed high-throughput mutagenesis assays for BTK function in B cells and T cells. We measured the fitness costs for thousands of point mutations in the PH–TH module and kinase domain, allowing us to assess whether dimerization of the PH–TH module and BTK kinase activity are necessary for function. In Ramos B cells we find that neither PH–TH dimerization nor kinase activity is required for BTK signaling. Instead, in Ramos cells, BTK signaling is enhanced by mutations in the PH–TH module that increase membrane adsorption, even at the cost of reduced PH–TH dimerization. In contrast, in Jurkat T cells, we find that BTK signaling depends on both PH–TH dimerization and kinase activity. Evolutionary analysis shows that BTK proteins in fish and lower organisms, like all Tec kinases other than BTK, lack PH–TH dimerization but have active kinase domains. Thus, PH–TH dimerization is not intrinsically required for Tec-kinase activity, and is a special feature that evolved to exert stricter regulatory control on BTK kinase activity as adaptive immune systems gained increased complexity.

## Introduction

One of the remarkable success stories in cancer treatment is the development of the small-molecule kinase inhibitor ibrutinib, which targets the Tec-family kinase BTK. BTK plays a critical role in B-cell receptor signaling and is dysregulated in certain leukemias [1]. In the canonical model, BTK phosphorylation at the membrane results in phosphorylation of phospholipase C γ (PLCγ), which generates the second messengers inositol trisphosphate and diacylglycerol, triggering calcium flux and initiating various signaling pathways [2]. The activation of PLCγ represents a transition from receptor-proximal to cell-wide signaling [3], and the activation of BTK is a gatekeeper of this transition.

BTK binds to phosphatidyl inositol lipids in the plasma membrane through its pleckstrin-homology and tec-homology domains (the PH–TH module) [4–6]. PH domains are phosphoinositide-binding domains that are present in at least ∼300 different human proteins [7]. PH domains typically bind to phosphoinositide as monomers and without strong specificity [8]. The PH domain of BTK binds with highest affinity to phosphatidylinositol (3,4,5)-trisphosphate (PIP_3_) [6], and unusually for PH domains, it can dimerize on membranes [4, 5]. The first evidence for dimerization came from a crystal structure of the BTK PH–TH module, determined by Saraste and colleagues, in which dimeric PH–TH molecules were observed in the crystal lattice [9] (Figure 1A). This dimeric arrangement, referred to here as the “Saraste dimer”, has been observed in all other crystal structures of the BTK PH–TH module that have been determined so far, whether alone or as part of larger constructs [4]. The PH–TH dimer in these structures has an interface composed of two loops: the S1 loop (connecting β strands 3 and 4 in the PH domain; residues 41–51 in human BTK), and the S2 loop (connecting β strands 5 and 6; residues 67–100) (Figure 1A) [9]. The S1 loop controls lipid specificity in the BTK PH domain [6], as it does in PH domains from other proteins [8].

**Figure 1.**
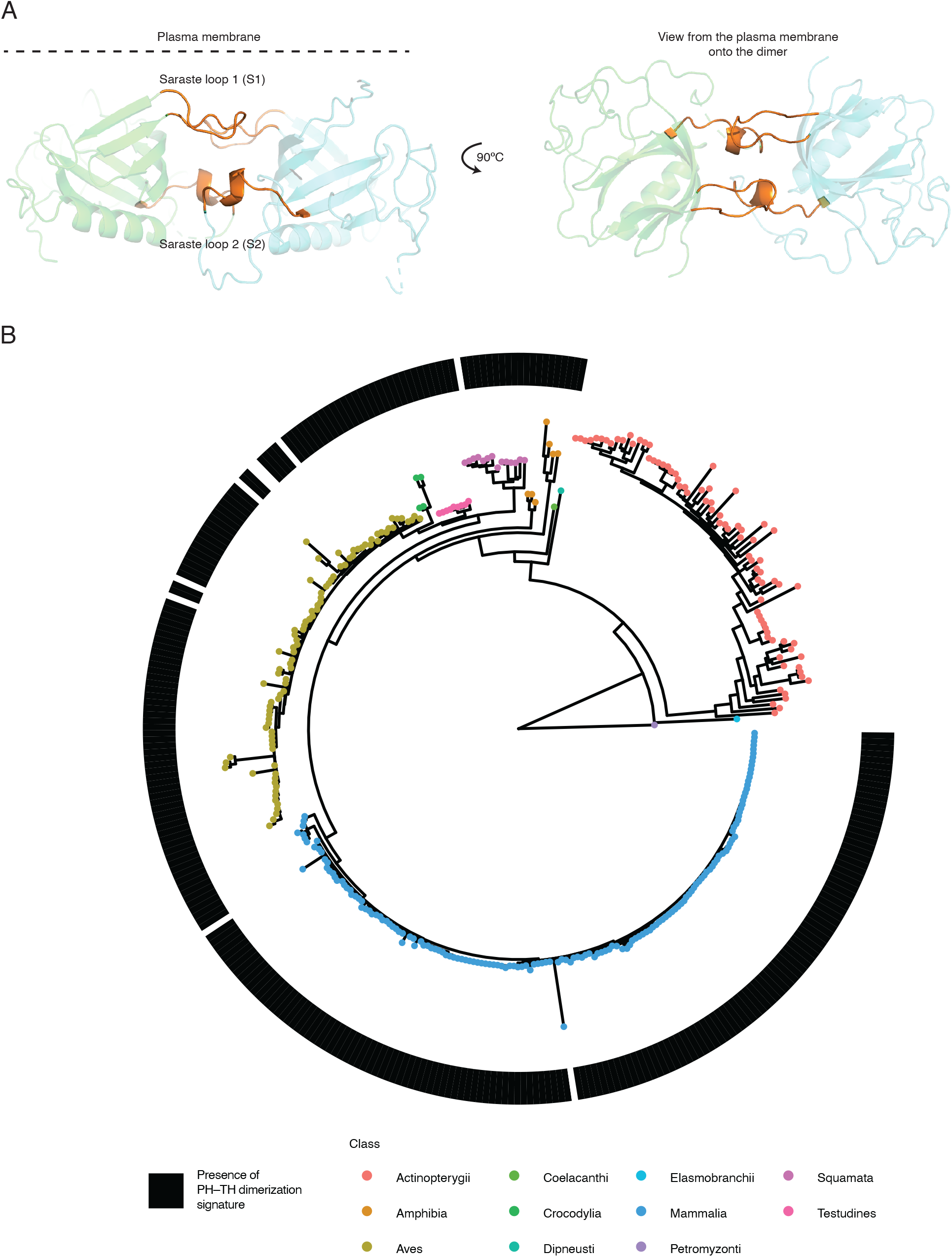
Evolution of the Saraste dimer interface. (A) Crystal structure of the Saraste dimer of the BTK PH–TH module (PDB: 1BTK). Loops S1 (residues 41–51) and part of S2 (residues 91–100) are shown in orange. Note that the insertion (residues 79–99, Figure 1B) includes the S2 loop and additional residues. (left) The membrane is shown as a dashed line. (right) The model is shown as if looking down through the membrane from the extracellular side. (B) A gene phylogenetic tree of BTK homologs. 366 sequences of BTK in representative vertebrates are shown. The tree is rooted using the lamprey (*Lethenteron camtschaticum*) Tec kinase. Vertebrate classes are shown as a single color. Homologs that have any sequence that aligns to residues 79–99 (IPRRGEESSEMEQISIIERFP) from human BTK (the dimerization signature) are denoted with a black rectangle (key). Others only have gaps in this region. Nine sequences outside of the Actinopterygii class are also missing the dimerization signature. These sequences are missing substantial N-terminal portions of the kinase as well and do not represent proteins that have lost the dimerization signature specifically.

The Saraste dimerization interface is composed of hydrophobic interactions made by residues presented by the S1 and S2 loops. The residues at the interface include Phe^44^ and Tyr^42^ (presented by the S1 loop) and Ile^92^ and Ile^95^ (presented by the S2 loop). Additional interactions between Leu^29^ and Ile^9^, presented by the neighboring β strands, also appear important for the interface. Mutations of these residues disrupt dimerization, as inferred from the observation that these mutations decrease BTK autophosphorylation [4]. These mutations also reduce the concentration dependence of the diffusion rate of BTK PH–TH molecules on supported-lipid bilayers, which is indicative of decreased dimerization [5]. Long time-scale molecular dynamics simulations show that these mutations induce immediate dissociation of the PH–TH dimer [10].

The ability of the BTK PH–TH module to dimerize on PIP_3_-containing membranes is intriguing because dimerization is a common mechanism by which tyrosine kinases are activated [11]. BTK dimerization through other mechanisms can enhance BTK activity. For example, artificial dimerization of a CD16–BTK fusion protein induces a rise in intracellular calcium [12]. The HIV protein Nef can also induce BTK dimerization through the SH2 and SH3 domains of BTK, resulting in activation [13]. A crystal structure of the kinase domain of BTK shows a kinase-domain dimer where the activation loop of one protomer is inserted into the active site of the other in a conformation reminiscent of trans autophosphorylation [14]. These observations suggest that the PIP_3_-dependent dimerization of the BTK PH–TH module is a mechanism to couple BTK activity to the levels of PIP_3_ in the plasma membrane.

The Tec-family kinases all have PH–TH modules (except for TXK) but the structural elements responsible for dimerization of the BTK PH–TH module are absent or drastically altered in other Tec-family kinases. The S1 loop, a key element of the PH–TH dimer interface in BTK, is shorter in other Tec kinases, and residues that contribute to the hydrophobic core in the BTK dimer are absent. Even for BTK, PH–TH dimers cannot be observed in solution, and seem to form exclusively on membranes. Dimerization of the BTK PH–TH module requires relatively high levels of PIP_3_ (∼1% of total membrane lipids) to occur at all [5]. Studies of lipid composition have described much lower total PIP_3_ levels (∼0.03% of total membrane lipids, rising to ∼0.4% upon cell stimulation [15]) but the local concentration of PIP_3_ can differ across the cell membrane. Whether the BTK PH–TH dimer is important for signaling in cells has not been established.

BTK has many functions in different cell types. In the classical view, the Tec kinase ITK is confined to T-cell lineages, and BTK, discovered because of its role in X-linked agammaglobulinemia [16], is expressed in B cells. But BTK has a wider range of expression, with roles in macrophages [17], CD4 and regulatory T cells [18], other myeloid lineages [19], and dendritic cells [20]. That BTK may be playing different roles in different cell types has been highlighted by studies showing that in some cells, BTK is required for signaling but its kinase activity is not [21–25]. In diffuse large B-cell lymphoma (DLBCL), inhibition of BTK kinase activity is an effective treatment, indicating that cancer progression requires kinase-active BTK. Unexpectedly, some ibrutinib-resistance mutations found in patients disrupt BTK kinase activity [22]. This suggests that there are important non-kinase functions for BTK. These function include the activation of phosphatidylinositol 4-phosphate 5-kinases (PIP5Ks) [24] and the activation of other kinases that are capable of phosphorylating PLCγ2, such as the Src-family kinase HCK [21]. Other studies have shown that the non-kinase functions of BTK are also important in normal physiology, since kinase-inactive BTK can rescue many aspects of B-cell development in a BTK-knockout mouse [26].

In this study, we set out to understand how BTK PH–TH dimerization contributes to signaling in lymphocytes, using Ramos B cells and Jurkat T cells as model systems. We developed high-throughput assays in these cells that report on how mutations affect BTK function, and use these assays to examine residues that modulate PH–TH dimerization. We interpret our results in light of the known structures of the PH–TH module [4, 9]. Unexpectedly, we find that BTK signaling depends on dimerization in the Jurkat but not the Ramos system, and that these differences reflect conditional requirements for BTK kinase activity in these two cell types.

## Results

### A conserved signature for BTK PH–TH dimerization

The BTK PH–TH module has a dimerization motif that is absent in the other Tec kinases. Both the S1 and S2 loops of the BTK PH–TH module were shown previously to be critical for dimerization [4, 5, 10], but the S2 loop differs from that of the other Tec kinases because it is present in a longer form only in BTK [10, 27]. In ITK, for example, this loop is 14 residues long, whereas in BTK it is 36 residues long. Only the longer form of the S2 loop can support dimerization of the PH–TH module, because the tip of the extended loop bears hydrophobic residues (Ile^92^ and Ile^95^) that are important for dimerization [4]. Thus, the longer S2 loop and its hydrophobic residues are a signature of BTK PH‒TH modules that are potentially capable of dimerization. Indeed, the ITK PH–TH module, with a shorter S2 loop, does not dimerize on membranes [5]. The S1 loop, by contrast, is conserved among Tec kinases. This loop is expected to be located proximal to the plasma membrane, and it presents charged residues that tune the affinity of BTK and other Tec kinases for membrane lipids [6].

We wondered when the signature for BTK PH–TH dimerization appeared during the evolution of BTK. We performed a phylogenetic analysis of BTK to provide an evolutionary context for the ability of the BTK PH–TH module to dimerize. Genes encoding Tec kinase proteins can be found in both jawed (Gnathostomata) and jawless (Agnatha) vertebrates even though adaptive immunity is thought to have arisen after the split between these two groups [28]. But genes encoding BTK proteins are first clearly identifiable only in the jawed-vertebrate lineage. Among extant jawed vertebrates, predicted BTK proteins can be distinguished from predicted ITK proteins because BTK proteins have higher pairwise identity to the human homolog. For example, predicted BTK proteins in 73 fish species have a median of 17% higher amino-acid identity to human BTK (median pairwise identity = 63%) than to human ITK (median pairwise identity = 46%).

The length of the S2 loop is not conserved in BTK proteins in jawed vertebrates [10, 27]. Most of the variation in length likely results from an exon insertion (residues 81–100) [27]. Furthermore, Ile^92^ and Ile^95^ are not conserved among fish, indicating that the signature for PH– TH dimerization is not conserved [10]. We constructed a BTK gene tree using representative sequences from different vertebrate classes (Figure 1B). This analysis used a multiple-sequence alignment of the full BTK sequence in 366 BTK proteins. The tree was rooted using the lamprey (*Lethenteron camtschaticum*) Tec kinase (the only non-BTK ortholog in the tree) [28].

We then overlaid the presence or absence of the extended S2-loop signature (residues 81– 100) onto the gene tree (Figure 1B). The result shows that the extended S2-loop signature, characteristic of human BTK and consistent with dimerization, arose after the split between ray-finned fishes (Actinopterygii), which do not have the signature, and lungfish (Dipneusti), which do. BTK proteins in the lobe-finned fishes (Coelacanth) also have the extended S2-loop signature, as do all of the other classes examined (Amphibia, Reptilia, Aves, and Mammalia). The hydrophobic residues Ile^92^ and Ile^95^ are conserved within these lineages as well. The S1 loop, important for phosphoinositide recognition in the membrane, is conserved among all these species, including the ray-finned fishes. These observations suggest that the original function of the BTK PH–TH module required membrane engagement but not PH–TH dimerization.

### High-throughput assays for BTK function in B cells and T cells

To understand how dimerization of the PH–TH module of BTK affects signaling in lymphocytes, we developed a high-throughput mutagenesis assay for BTK function. This assay relies on monitoring the cell-surface expression of CD69, a C-type lectin protein that is upregulated in response to several different types of stimulation [29]. We tested whether BTK can increase CD69 expression in Ramos B cells and Jurkat T cells. Ramos cells are a Burkitt’s Lymphoma B-cell line, derived from a human male, and these cells therefore contain only one copy of the *BTK* gene [30]. Unlike many other B-cell lines, the Ramos line does not depend on BTK for cell growth [23]. Nevertheless, Ramos cells exhibit BTK-dependent calcium responses [23] and correlated CD69 upregulation [31].

Jurkat T cells are a T-cell leukemia line derived from a human male [32]. Heterologous expression of BTK in Jurkat T cells has been used previously to study BTK function, again by monitoring calcium responses and CD69 upregulation [33, 34]. Compared to Ramos cells, Jurkat cells have a related but distinct complement of adaptor proteins. In Jurkat cells and T cells more generally, SLP76 in combination with Linker for Activated T-cell (LAT) and GRB2-related adaptor protein (GADS) recruits ITK to the signalosome. LAT is not expressed in B cells. Instead, B cells express SLP65, which shares substantial sequence and structural homology with SLP76, but seems to combine the functions of LAT, SLP76, and GADS in one protein [35, 36]. In addition to these differences in adaptor proteins, Jurkat cells have elevated levels of PIP_3_ due to loss of PTEN function [37, 38]. Loss of PTEN function is important in some cancers, including some diffuse large B-cell lymphomas [39].

We developed an assay for BTK function by first knocking out the single allele of BTK in Ramos cells using CRISPR/Cas9 (Figure S1A). We then added back human BTK with a lentivirus, using an internal ribosome entry site (IRES) followed by EGFP to monitor lentiviral transduction. To assess stimulation, we monitored CD69 expression using FACS in the presence or absence of the anti-IgM (µ chain) stimulatory antibody.

As expected, removal of BTK from the Ramos cells dampened CD69 responses overall (Figure 2A–B). In the knockout without stimulation, there is complete absence of basal CD69 expression. Re-addition of BTK to the knockout cells induces CD69 expression in the presence of stimulation, indicating that BTK activity still requires the endogenous regulatory signals when the protein is added using a lentivirus (Figure 2B). In keeping with previous reports, some CD69 expression is observed upon stimulation of the knockout cells even without BTK, indicating that other signaling molecules or pathways contribute to the B-cell response in these cells [23]. Even in wild-type Ramos cells, some CD69 expression is observed without stimulation or additional expression of a kinase (Figure 2A), presumably due to the low but detectable amount of phospo-PLCγ2 constitutively present in this cell line [40]. This is further underscored by many examples of seeming redundancy in B-cell signaling pathways [41].

**Figure 2.**
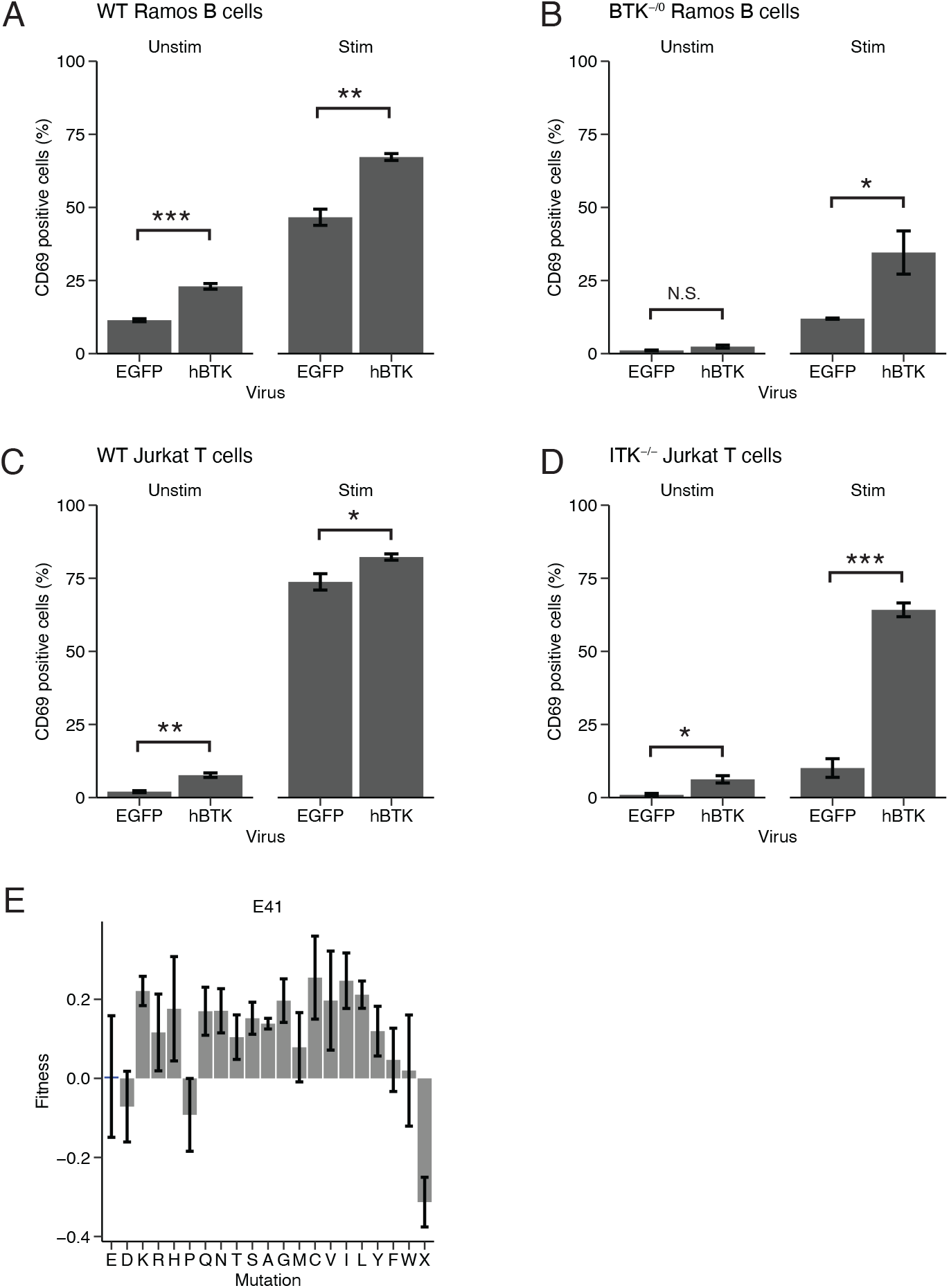
A high-through mutagenesis assay for BTK function in B cells and T cells. (A) BTK induces CD69 expression in Ramos cells. The percentage of the population that expresses CD69, measured by FACS, is shown. Cell populations were transduced four days before sorting with lentiviruses expressing EGFP or human BTK. The panel at left is without stimulation. At right, the cells were stimulated with anti-IgM (µ chain) for 16 h at 10 µg/mL. Shown is the mean ± standard error from three replicates. N.S: not significant, *p < 0.05, **p < 0.01, ***p < 0.001, t test. (B) BTK-deficient Ramos cells induce CD69 expression in response to stimulation, and exhibit increased CD69 upon expression of human BTK without stimulation. Otherwise as in (A). (C) BTK induces CD69 expression in Jurkat cells. The cells were left unstimulated (left) or stimulated (right) with 1 µg/mL anti-T cell receptor antibody C305 for 16 h. Shown is the mean ± standard error from four replicates. (D) ITK-deficient Jurkat cells induce CD69 expression in response to stimulation, and exhibit increased CD69 upon expression of human BTK without stimulation. Otherwise as in (A). (E) The effects of mutations at Glu^41^ agree with previous studies. Mean fitness values ± standard error for biological triplicates are shown (y axis) for all possible amino acid substitutions (x axis). X represents the average value for two stop codons. The wild-type residue is shown in blue, with the residue single-letter code and human numbering shown above the block of substitutions.

In Jurkat cells, BTK induces strong CD69 expression in response to stimulation with the T-cell receptor (TCR) crosslinking antibody C305 [42] (Figure 2C–D). The effect of stimulation is even more apparent when using Jurkat cells in which ITK has been knocked out because the basal levels of CD69 expression in these cells is lower (Figure 2D). BTK expression also induces a modest but detectable amount of CD69 expression in the absence of stimulation, both in ITK-deficient and in wild-type Jurkat cells (Figure 2C–D). Addition of ibrutinib decreased BTK-induced CD69 upregulation in the absence (Figure S1B) and presence (Figure S1C) of stimulation, indicating that BTK activity is required for both the basal and stimulation-induced CD69 upregulation. These observations indicate that BTK can compensate for the signaling deficit induced by ITK loss in Jurkat cells.

These observations in Ramos and Jurkat cells guided the construction of high-throughput mutagenesis assays for BTK function. Lentiviruses were prepared with sequences encoding wild-type BTK or variants, and used to transduce a pool of BTK-deficient Ramos or ITK-deficient Jurkat cells at low multiplicity of infection (Figure S1D). CD69-expressing cells were separated from non-expressers using FACS or magnetic-bead separation (see methods). mRNAs derived from BTK genes in both the input population and the selected population were sequenced, and the resulting counts from each variant in both populations were used to generate fitness scores by comparing to the wild-type population.

We used mRNA-seq to measure the enrichment of each BTK variant in a pool of cells enriched for high CD69 expression. Most previous high-throughput mutagenesis studies have sequenced genomic DNA (referred to as gDNA) to quantify the relative population of variants [43–45]. But in our experiments, mRNA-seq, used to measure mRNA levels, yielded consistently lower variability between replicates than gDNA-seq. This is presumably due to both the increased copy numbers of mRNA compared to gDNA in each cell and the amplification that occurs in the reverse transcription step where cDNA is made from mRNA. During this step, cDNA is produced iteratively in a process that results in linear amplification of the mRNA sequence, which maintains copy numbers more precisely than the exponential PCR amplification of gDNA [46]. The use of mRNA-seq to count variant levels has two additional advantages: (1) because the reverse-transcription step uses a primer that anneals to the IRES, only mRNAs from the viral construct are sequenced, avoiding any endogenous transcripts. (2) mRNA copy number is a better correlate of protein expression than gDNA copy number because some mutations alter mRNA copy number but leave gDNA copy number unchanged. For example, mutations that cause nonsense-mediated decay decrease the stability of mRNAs [47]. We see evidence for nonsense-mediated decay (either in the Ramos cells or in the lentiviral-packaging HEK293FT cells), as mRNAs with stop-codon mutations are consistently among the least well represented in our unselected libraries compared to mRNAs with other mutations (Figure S1E–F).

mRNA-seq read counts in the CD69-selected libraries were normalized to the unselected pool and to the wild-type sequences to develop a metric for “fitness”, according the equation 1:

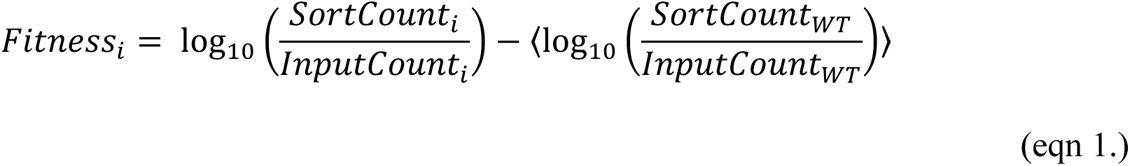

Here, *Fitness_i_*, *SortCount_i_* and *InputCount_i_* are the fitness score, read counts in the CD69-sorted library, and read counts in the input library for a particular variant *i*, respectively. *SortCount_WT_* and *InputCount_WT_* are the average read counts for the 20–30 different sequences encoding wild-type proteins included in each library. Fitness values above zero indicate that a particular BTK variant induces more CD69 expression than wild-type BTK (gain-of-function alleles), while values below zero correspond to variants that show decreased CD69 expression relative to wild-type (loss-of-function alleles). At some positions, we observe that synonymous codon substitutions of the wild-type codon change fitness slightly. This may be attributed to synonymous codon effects on translation rate [48], and in our datasets these effects are small.

To determine the optimal concentration of stimulatory antibody with which to measure BTK fitness scores, we performed a concentration-course of stimulatory antibody and analyzed the resulting mRNA-seq data. We stimulated ITK-deficient Jurkat cells with 8 concentrations of TCR crosslinking antibody and prepared 16 mRNA-seq libraries of mutations in the S1 loop: 8 for the input and 8 for the CD69-sorted populations (Figure S2A). As the stimulant concentration is increased, almost all positions in the library tolerate more variation. This can be observed by the increasing overlap of fitness scores for the mutants and the wild-type proteins (Figure S2B). It can also be observed by the overall weakening of the deleterious effects for some mutants as the stimulatory antibody concentration is increased (Figure S2C). We performed a similar experiment in the BTK-deficient Ramos cells, stimulating the cells with 4 concentrations of anti-IgM stimulatory antibody. In this experiment, the range of fitness scores also decreases with increasing antibody concentration, as in the Jurkat system. Because of this, the biological-replicate agreement, measured by the Pearson correlation between fitness scores from different Ramos samples, decreases with increasing stimulatory antibody concentration (Figure S2D). Overall, these experiments in both the Jurkat and Ramos systems showed that the best measurements of relative fitness were obtained at the lowest concentrations of stimulatory antibodies. Presumably, high levels of stimulation in both the Ramos and Jurkat cells can bypass the requirement for BTK in CD69 upregulation, leading to increased noise in the data.

To assess our approach, we examined the well-characterized BTK mutation E41K in Ramos cells (Figure 2E). E41K was previously identified as a gain-of-function mutation in BTK that promotes non-specific membrane binding by the PH–TH module [6, 49, 50]. Consistent with this, we observe that the E41K BTK variant significantly increases CD69 expression in Ramos cells (fitness score = 0.22 ± 0.04, Figure 2E). One explanation for the activating effect of the E41K mutation is the removal of a negatively charged residue (Glu^41^) from the surface of the PH domain that interacts with the membrane. A second explanation, which is not mutually exclusive with the first, is that E41K is activating because of the addition of a positively charged residue (lysine) to the membrane-interacting surface of the PH domain. We find that many substitutions of Glu^41^ confer a fitness advantage, not just the lysine, consistent with the principal activating effect being the loss of the negatively charged residue, rather than addition of the positively charged one. Recent coarse-grained molecular dynamics simulations of the BTK PH domain support this interpretation [51]. Not all substitutions at position 41 confer a fitness advantage, however. Proline distorts the polypeptide backbone, and in our data the mutation E41P (–0.09 ± 0.09) is neutral or slightly loss-of-function. Substitution by aspartate, which is chemically similar to glutamate, is neutral (–0.07 ± 0.08).

### PH–TH dimerization is dispensable for BTK signaling in Ramos B cells

In order to understand how PH–TH dimerization influences BTK signaling, we generated two libraries of variants in which mutations span the S1 loop (residues 41–51) and part of the S2 loop (residues 91–100). In these libraries, every residue was changed to ever other possible amino acid and stop codons were introduced at each position. The libraries were designed with two codons per residue were (except for the methionine and tryptophan residues), two stop codons, and several redundant versions of the wildtype sequence that each use 5 alternate codons. This redundancy was key to correlating the fitness scores to amino-acid substitutions rather than codon substitutions. In total, the S1 library was composed of 443 sequences and the S2 library was composed of 401 sequences. The S1 and S2 regions were chosen because the interpretation of the mutations from these libraries could be anchored on previous studies that characterized the roles of these loops in BTK dimerization [4, 5, 10].

The key hydrophobic sidechains at the Saraste-dimer interface include Tyr^42^, Phe^44^, Ile^92^, and Ile^95^ [4] (Figure 3A). Three previous studies [4, 5, 10] have used mutations at Tyr^42^ to investigate dimerization in experiments using purified proteins, and consistent with their results, we find that all substitutions to Tyr^42^ are loss-of-function (Figure 3B). To our surprise, however, mutation of other hydrophobic residues at the dimer interface did not lead to loss of function.

**Figure 3.**
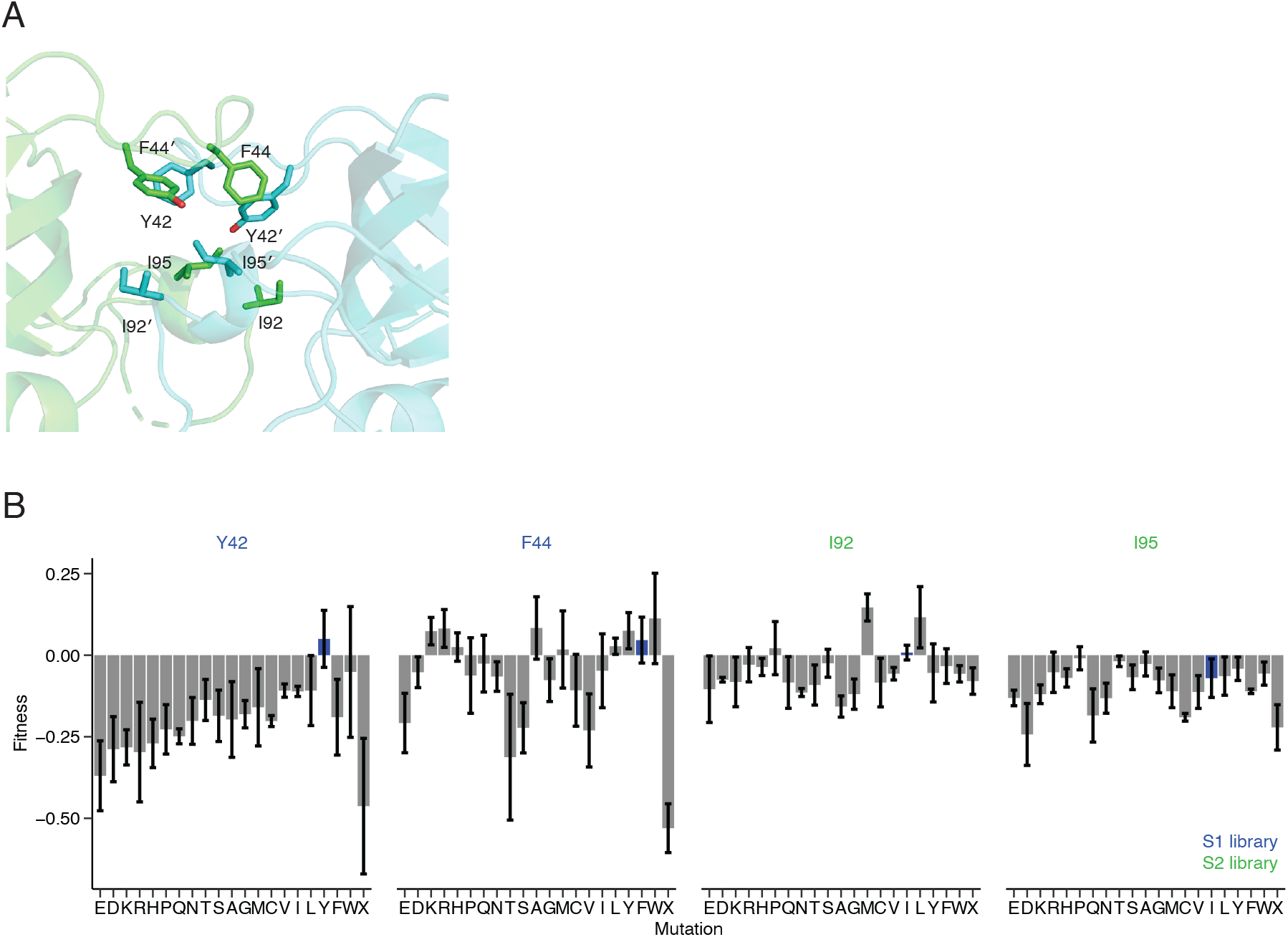
PH–TH dimerization is dispensable in Ramos B cells. (A) The hydrophobic residues that constitute the Saraste dimer interface. PH‒TH dimer interface residues are shown as sticks on a cartoon representation of the dimer interface, with the two protomers shown in different colors (PDB: 1BTK). (B) Sensitivity of the PH‒TH dimer interface mutations in Ramos cells. For each residue shown in (A), fitness scores for all possible mutations at that position are shown. Tyr^42^ and Phe^44^ values are from the S1-loop library. Ile^92^ and Ile^95^ are from the S2-loop library (key). Otherwise as in Figure 2E.

The sidechain of Phe^44^ in one protomer packs against the side chain of the same residue in the other protomer and we would expect mutations in Phe^44^ to disrupt the dimer interface. However, substitution of Phe^44^ by lysine (fitness score = 0.07 ± 0.04) or arginine (0.08 ± 0.06) increases, rather than decreases, fitness. Ile^95^ also forms part of the dimer interface, packing against Ile^95^ and Tyr^42^ in the other protomer and stabilizing Phe^44^ in the same protomer. Substitutions of this residue with charged residues would be expect to disrupt dimerization. But substitution of Ile^95^ by arginine (–0.05 ± 0.06) or proline (–0.01 ± 0.04) are both tolerated. The proline in particular is expected to disrupt the small helix presented by the S2 loop, reconfiguring its reciprocal presentation of hydrophobic sidechains at the interface. Overall, these data indicate that PH–TH dimerization is not critical for BTK signaling in Ramos B cells. As discussed below, Tyr^42^ seems to be important for the stability of the PH domain, and the loss of function due to mutations of this residue may be a consequence of destabilization of BTK.

### Ramos-cell signaling does not depend on BTK kinase activity

The role of BTK in some human B-cell lines, and the Ramos line in particular, is unusual. BTK variants with mutations that disrupt BTK kinase activity fully support BTK-induced calcium flux in Ramos and TMD8 cells [21, 23–25]. These results are reminiscent of studies that show that expression of a *Btk* allele with a K430R mutation in the kinase domain (which alters the interaction of the kinase with ATP and attenuates or abolishes kinase activity [52]) does not prevent BTK from supporting most of B-cell development in a *Btk*-knockout mouse [26]. They are also consistent with a growing body of literature that describes non-catalytic roles for BTK in a number of cancers, including DLBCL [22]. Because the Src, Syk, and Tec families of protein tyrosine kinases can all phosphorylate Plcγ2 in vitro [40], it seems that BTK in some contexts may function more as a scaffold than as a kinase, acting to support the proper localization or activation of other kinases.

We wondered whether BTK PH–TH dimerization is only important when BTK kinase activity is important. To study this, we mutated residues that are important for BTK kinase activity, and scored the effects of the mutations using our assay. We targeted the histidine– arginine–aspartate (HRD) motif in BTK (residues 519–521) (Figure S3A). This motif is highly conserved among human tyrosine kinases (present in 90% of sequences), with the aspartate residue in the motif being essentially invariant. Among the vertebrate Tec kinases, the HRD motif is present in 98% of sequences, with the aspartate present in 99%. The aspartate residue in the HRD motif is located below the terminal phosphate of ATP and the hydroxl group of the substrate tyrosine. It is referred to as the “catalytic base” because it is likely to abstract a proton from the substrate hydroxyl group [53]. Mutation of the HRD aspartate abolishes kinase activity in other tyrosine kinases [54]. We constructed a library of point mutations at positions including and surrounding the HRD motif, including two codons per residue (except for the methionine and tryptophan residues), two stop codons, and several redundant versions of the wildtype sequence. In total, the library comprised 500 sequences and spanned 12 positions in the BTK kinase domain.

The absolute values of the fitness scores from the HRD library were overall much smaller than for the S1 and S2 loop libraries, and the fitness scores showed more variability (Figure S3B). The decreased magnitude of the scores indicated that the motif exhibits substantial tolerance for substitution, which is unexpected given its central location at the heart of the catalytic center of the kinase domain. Consistent with this, His^519^ tolerates substitution by both lysine (fitness score = 0.01 ± 0.01) and isoleucine (–0.01 ± 0.01). Strikingly, Asp^521^ in the motif tolerates substitution by phenylalanine (0.00 ± 0.01) and alanine (–0.01 ± 0.01). Substitution of Asp^521^ is not compatible with kinase activity: indeed, mutation of the corresponding aspartate to alanine in yeast PKA reduces the activity of the kinase to 0.4% of wild-type [55]. The overall insensitivity of CD69 expression levels to mutations in this region indicates that BTK signaling in Ramos cells does not require BTK kinase activity.

### Kinase activity is required for BTK to substitute for ITK in T cells

We investigated the HRD-motif library using ITK-deficient Jurkat cells instead of BTK-deficient Ramos cells. In stark contrast to our results from the Ramos system, the fitness scores from these experiments exhibited low variability and strong signals (Figure 4A and Figure S4). Examining the effects of substituting the catalytic base, Asp^521^, we find that it tolerates no substitutions (Figure 4A). All mutations at this position decrease BTK function to the same extent as a stop codon (X), indicating that the ability of BTK to induce CD69 expression in Jurkat cells depends on BTK kinase activity. At other positions, even conservative substitutions lead to substantial fitness loss, even if their effects are not as severe as the stop-codon substitution. For example, substitution of Arg^520^ by lysine (Figure 4A) results in substantial fitness loss (fitness score = – 0.24 ± 0.03, compared to –0.39 ± 0.01 for the stop codons). Similarly, L518I is also a loss-of-function mutation (–0.16 ± 0.03). This hypersensitivity of residues in the HRD motif and in surrounding regions (Figure S5A) is consistent with the high conservation of these residues and their critical roles in kinase activity.

**Figure 4.**
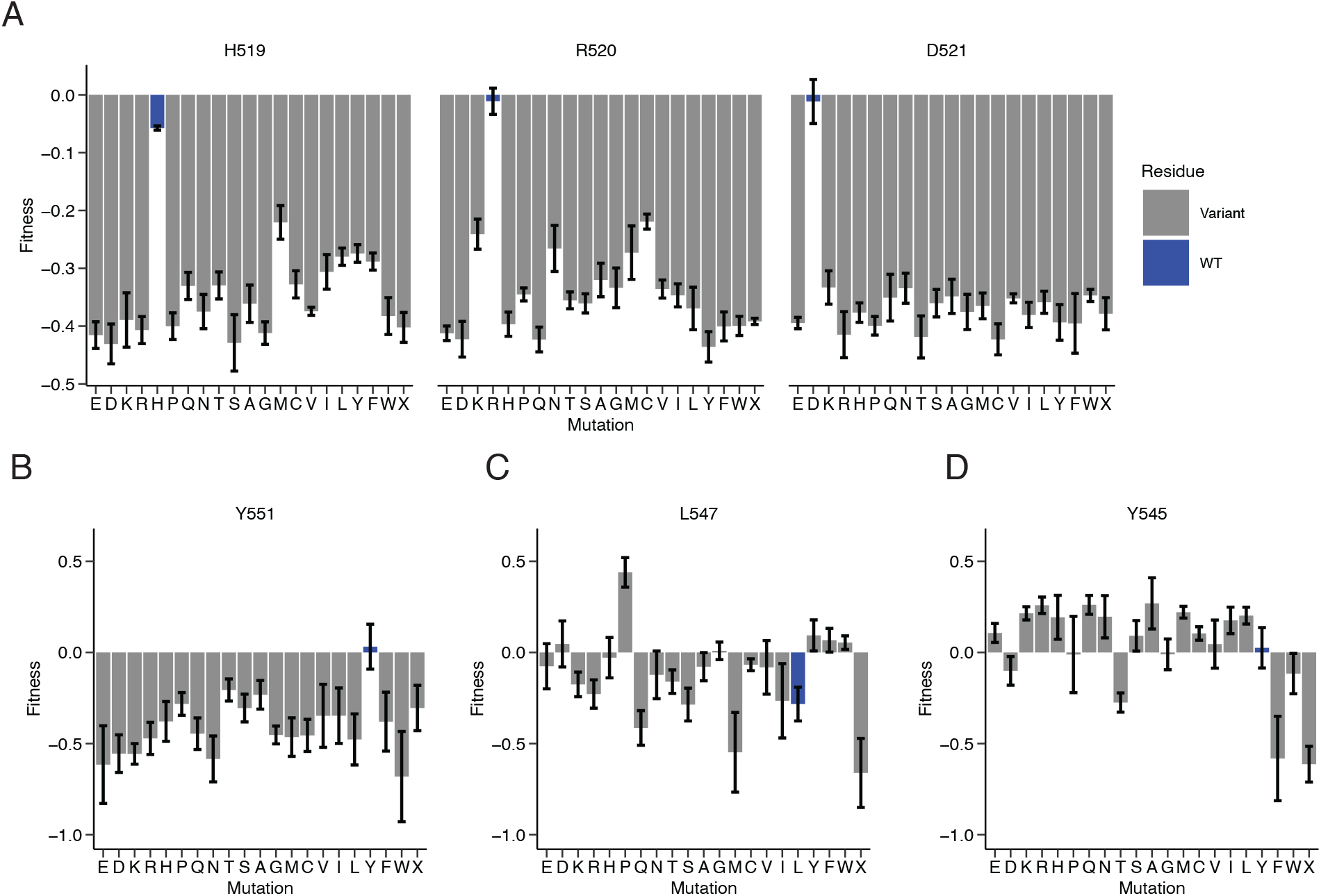
Kinase activity is required for BTK signaling in Jurkat T cells. (A) The HRD motif is sensitive to mutation in Jurkat cells. Fitness scores for all possible substitutions in the HRD motif are shown, with the residue code and number above each block of mutations. N = 4 biological replicates. Otherwise as in Figure 2E. (B) Mutations in the phospho-regulated tyrosine Tyr^551^ in the activation-loop are not tolerated. Otherwise as in (A), except plotting values from the activation-loop library. (C) Leu^547^, when mutated to proline, is activating. Otherwise as in (B). (D) Tyr^545^, at the equivalent position as Val^600^ in BRAF kinase, increases BTK activity when mutated to many residues. Otherwise as in (B).

We also tested the effects of mutations in another part of the kinase domain that is critical for activity: the activation loop. The activation loop is a flexible segment of ∼20–25 residues within the C-lobe that controls kinase activation [56]. Mutagenizing this region and three flanking residues (positions 538-565, with the variation encoded in 1155 total sequences) showed loss and gain-of-function variants that strongly indicate that BTK kinase activity is required for signaling (Figure S5B). Tyr^551^ corresponds to the site of activating autophosphorylation in the activation loops of most tyrosine kinases. Phosphorylation of Tyr^551^ in BTK relies on either SYK or LYN in BCR signaling [57] and all mutations at this position result in fitness loss (Figure 4B). As expected, glutamate or aspartate at this position fail to mimic a phosphotyrosine residue. The L547P mutation (fitness = 0.44 ± 0.08, Figure 4C) is an unusual case where a proline substitution causes gain-of-function. This residue has been characterized previously as a mutation that confers resistance to ibrutinib [58]. Ibrutinib recognizes an inactive conformation of the kinase domain that is likely to be destabilized by the proline, leading to drug resistance and activation [59]. The experiments reported here were performed without inhibitors present, and so the fact that the proline mutation is strongly activating in this context is consistent with the mutation biasing the kinase domain toward the active conformation.

We examined the effect of mutations to Tyr^545^, which is at the equivalent position as Val^600^ in BRAF kinase (Figure 4D). The BRAF kinase domain adopts an inactive conformation that resembles that of BTK. Val^600^ makes hydrophobic interactions in the inactive state which are destabilized by the oncogenic BRAF mutation V600E [60]. In BTK, we find that mutation of Tyr^545^ to almost any charged or hydrophilic residues is activating. By contrast, hydrophobic substitutions of this residue to phenylalanine (–0.58 ± 0.23), tryptophan (–0.12 ± 0.11), glycine (–0.01 ± 0.08), or valine (0.05 ± 0.13), are neutral or loss of function. These data are consistent with a hydrophobic stabilizing interaction made by Tyr^545^ in a manner similar to that seen in BRAF, as expected from the structures of these two kinase domains. Collectively, the effects of mutations in the activation loop show that the kinase domain needs to convert to the active conformation for BTK signaling. When considered alongside the mutations in the HRD motif, these datasets indicate that kinase activity is required for BTK to substitute for ITK in Jurkat cells.

### BTK requires PH–TH dimerization in Jurkat cells

Given the strong dependence of the ITK-deficient Jurkat cells on BTK kinase activity, we asked whether PH–TH dimerization might be more important for signaling in this context compared to the Ramos cells. We transduced the ITK-deficient Jurkat cells with BTK variants encoded by the S1 and S2-loop libraries and monitored their ability to increase CD69 expression, as before. For the majority of residues, the resulting fitness scores from these two libraries agree well between the Ramos and Jurkat cell lines (Figure 5A and Figures S6A–B), with an R of 0.7 over the 441 fitness scores for each mutation at each position in these loops and a close adherence to the y = x line. This result indicates that most of our observations concerning how different substitutions in the PH–TH module affect BTK signaling do not depend on the model system. In particular, recognition of phosphoinositides by the PH domain is important in both Ramos cells and Jurkat cells.

**Figure 5.**
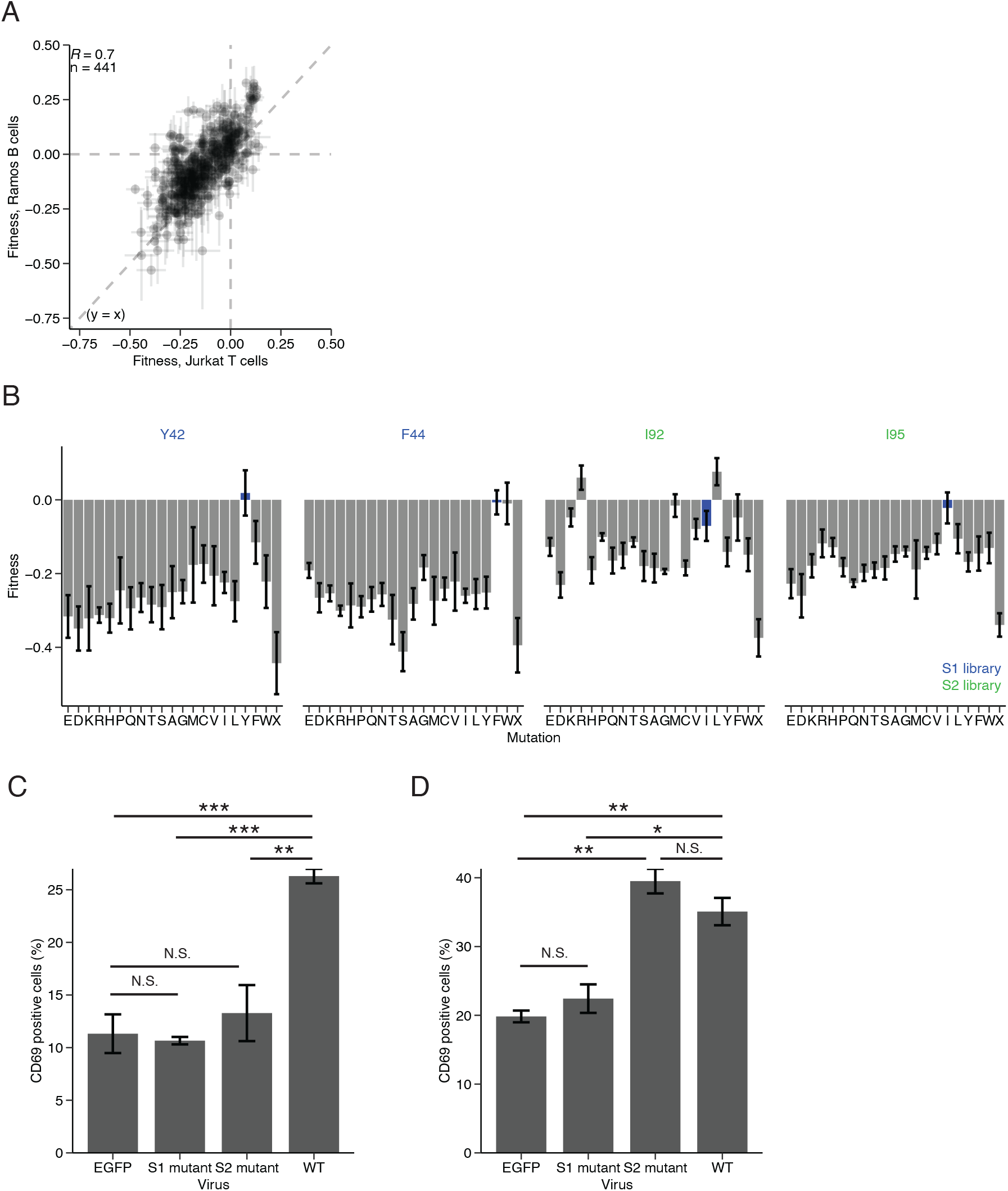
PH–TH dimerization is required for signaling in Jurkat T cells. (A) Agreement between fitness scores between the Ramos B cell and Jurkat T cell for the S1 and S2-loop libraries. Mean fitness scores ± standard error for n = 441 substitutions in the two libraries are plotted for both cell lines. Dashed lines indicate x = 0, y = 0, and y = x. (B) The dimer interface is sensitive in Jurkat cells. Otherwise as in Figure 3B, except showing data for Jurkat cells with n = 4 biological replicates. (C) Both the S1 and S2 loops are required for signaling in Jurkat T cells. Lentiviruses expressing human BTK without an abridged S1 loop (with residues 41–51 deleted, “S1 mutant”) or S2 loop (with residues 91–100 deleted, “S2 mutant”) were used to transduced ITK-deficient Jurkat T cells. The mean fraction of CD69-positive cells ± standard error, determined by FACS, is shown for n = 6 replicates. N.S: not significant, *p < 0.05, **p < 0.01, ***p < 0.001, t test. (D) The S2 loop is dispensable in Ramos B cells. Otherwise as in (E), except for n = 3 replicates.

Closer inspection of these data reveals key differences between the cell lines regarding the importance of PH–TH dimerization. For the Jurkat cell experiments, we find that the residues in the PH–TH dimer interface (Figure 3A) tolerate almost no substitutions (Figure 5B). For example, Phe^44^, the key hydrophobic interface residue, is very sensitive to mutation in Jurkat cells (Figure 5B), unlike in Ramos cells (Figure 3B). In Jurkat cells, it tolerates only substitution by tryptophan (fitness score = –0.01 ± 0.06). Similarly, Ile^95^ tolerates no substitutions in Jurkat cells but tolerates many substitutions in Ramos cells. Tyr^42^, the residue mutated to disrupt dimerization in previous studies [4, 5] tolerates no substitutions in either cellular context.

The sensitivity of Tyr^42^ to mutation in both cell lines may reflect a role for this residue in the stability of the PH domain, in addition to a role in dimerization. In contrast to Phe^44^, which makes no intramolecular sidechain contacts, the sidechain of Tyr^42^ packs against that of Ile^9^ and Leu^29^ from the same protomer, and these intramolecular contacts may be important for folding of the PH domain. Two previous studies have used mutations of Tyr^42^ to examine PH–TH dimerization in biochemical assays and chicken DT40 B cells [4, 5]. But the data presented here suggest that mutations to Tyr^42^ destabilize the PH domain, which would decrease the calcium flux in DT40 cells regardless of whether mutations to Tyr^42^ affect PH–TH dimerization.

To understand more about the relative importance of PH–TH dimerization in the Jurkat and Ramos contexts, we prepared constructs in which we deleted the entire S1 loop region (a membrane interacting loop; residues 41–51) or the exon insertion in the S2 loop (important for dimerization; residues 81–100). In the Jurkat cells, removal of either the S1 loop or the S2-loop insertion decreased CD69 expression to baseline levels. This supports the conclusion that both membrane binding and dimerization are required for signaling by BTK in Jurkat cells (Figure 5C). In Ramos cells, the protein without the S1 loop also fails to signal (Figure 5D) and is indistinguishable from the control virus. But BTK without the S2-loop insertion has no signaling deficit in Ramos cells. These data, combined with the saturation-mutagenesis data, support a role for PH–TH dimerization in Jurkat cells but not in Ramos cells.

### Membrane binding through the canonical lipid-binding site is required for signaling

The PH domain of BTK shares a canonical lipid-binding site with PH domains from other proteins. Biochemical and biophysical studies of the BTK PH–TH module have revealed another lipid-binding site, the peripheral site, that is occupied when the PH module forms a dimer [4, 5, 10]. To examine the sensitivity of these two sites in our assays, we generated two additional saturation-mutagenesis libraries that span the canonical lipid-binding site (residues 8–36, 1163 sequences) and the peripheral lipid-binding site (residues 52–62, 460 sequences). We used the Jurkat cells to measure fitness scores. As expected, the residues that constitute the canonical lipid binding site are sensitive to mutation (Figures 6A–B and S7A–D). In particular, Lys^12^, Asn^24^, Lys^26^, Arg^28^, and Lys^53^, which are important for recognition of the PIP_3_ headgroup [4], tolerate no substitutions without a loss in fitness.

**Figure 6.**
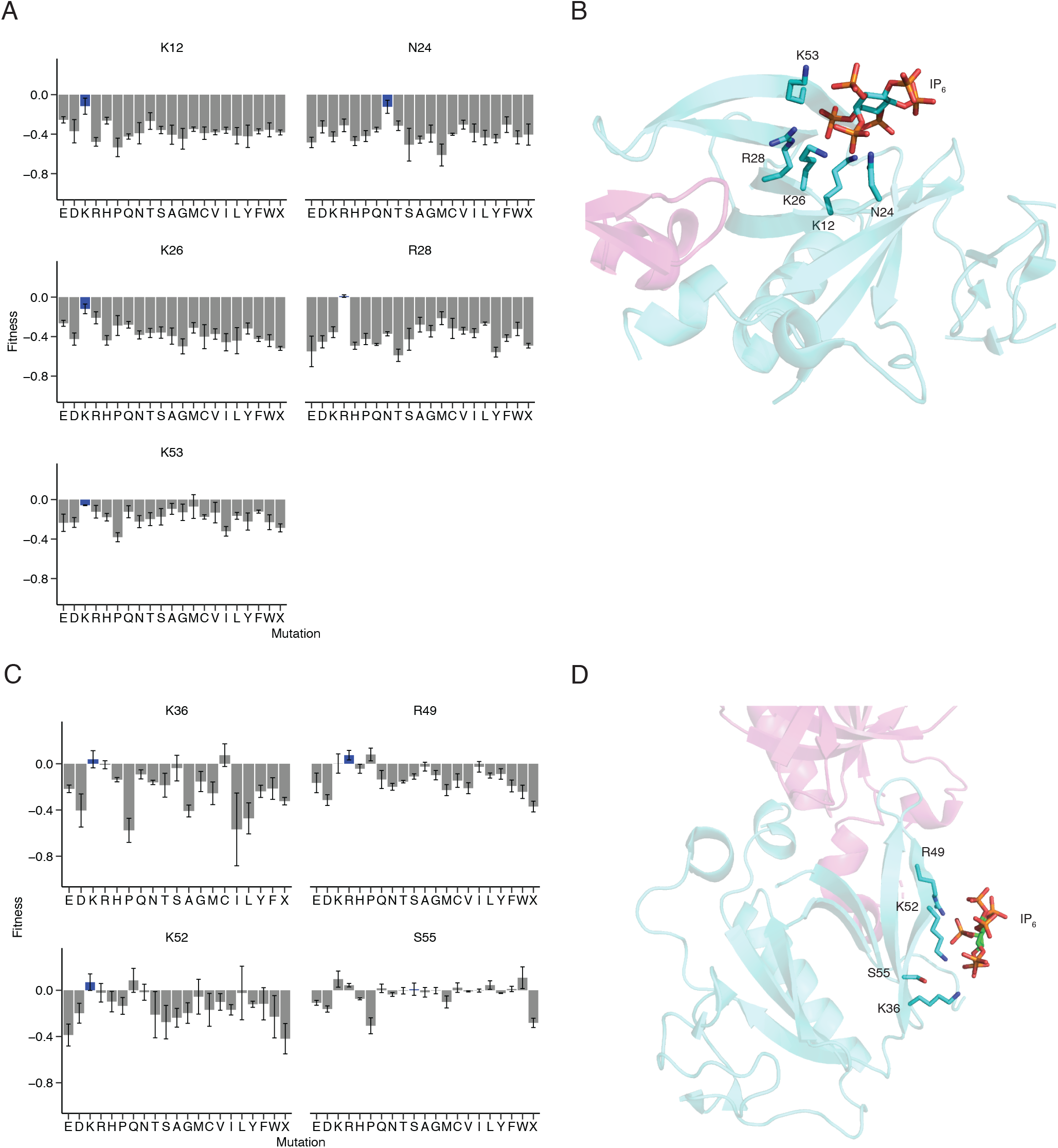
The canonical PIP_3_-binding site is sensitive to mutation but the peripheral site is tolerant. (A) The residues that constitute the canonical site are sensitive to mutation. Fitness scores for all substitutions for residues that interacts with the lipid are shown. Otherwise as in Figure 2E. (B) The crystal structure (PDB: 4Y94) of the PH–TH module of BTK is shown as a cartoon. The two protomers of the dimer are shown in cyan and pink, with the canonical-site interacting residues shown as stick representation on one of the two protomers. The lipid crystallized in this structure, hexakisphosphate (IP_6_) is shown as stick representation. (C) The residues of the peripheral site tolerate mutation. Fitness scores for all possible substitutions for each residue that interacts with the peripheral-site lipid are shown. Otherwise as in (A). (D) Crystal structure of the PH–TH module of BTK with the peripheral-site residues shown as stick representation. Otherwise as in (B).

The mutagenesis data show that the peripheral lipid-binding site in the BTK PH–TH module is not as critical for CD69 expression, as it tolerates more mutations. Consistent with a previous report [5], the R49S mutation (fitness score = –0.11 ± 0.02) and the K52S mutation (– 0.28 ± 0.14) both individually reduce signaling (Figure 6C–D). However, Lys^52^ tolerates substitution by arginine (–0.02 ± 0.08), asparagine (–0.02 ± 0.07), glutamine (0.09 ± 0.1), isoleucine (–0.02 ± 0.23), and methionine (–0.06 ± 0.15). Another residue at the peripheral site, Lys^36^, tolerates substitution by arginine (–0.01 ± 0.03), serine (–0.04 ± 0.1), and cysteine (0.07 ± 0.1). The tolerance at these sites suggests that PIP_3_ does not necessarily have to be bound at the peripheral site during signaling, despite the requirement for dimerization in these cells. Two modest gain-of-function substitutions, S55K (0.1 ± 0.07) and S55R (0.04 ± 0.02), indicate that more positive charge near this lipid site can enhance signaling (Figure 6C–D). These substitutions show that while the canonical lipid-binding site is crucial, the peripheral site can contribute to signaling, but is not required. Results from molecular-dynamics simulations [10] have suggested that the BTK PH–TH dimer engages multiple PIP_3_ molecules in addition to the one bound at the canonical site. Coarse-grain simulations of the Brag2 PH domain show that this domain can also engage multiple phosphoinositides [61]. Thus, single mutations at the peripheral site of the BTK PH–TH module may be less impactful.

### Context-dependent PH-domain mutations

In examining the fitness landscapes for the S1 and S2 libraries in the Ramos and Jurkat cells, we noticed a set of mutations that decrease fitness in Jurkat cells but increase fitness in Ramos cells. Surprisingly, many of these mutations are substitutions of neutral or negatively charged residues by lysine or arginine (Figure 7A). Glu^41^, Phe^44^, Glu^45^, Gly^47^, and Gly^50^, all of which are in the S1 loop, exhibit this switching behavior most strongly. By contrast, when the wild-type residue is positively charged (Arg^46^, Arg^48^, Arg^49^), it does not exhibit differential fitness when mutated.

**Figure 7.**
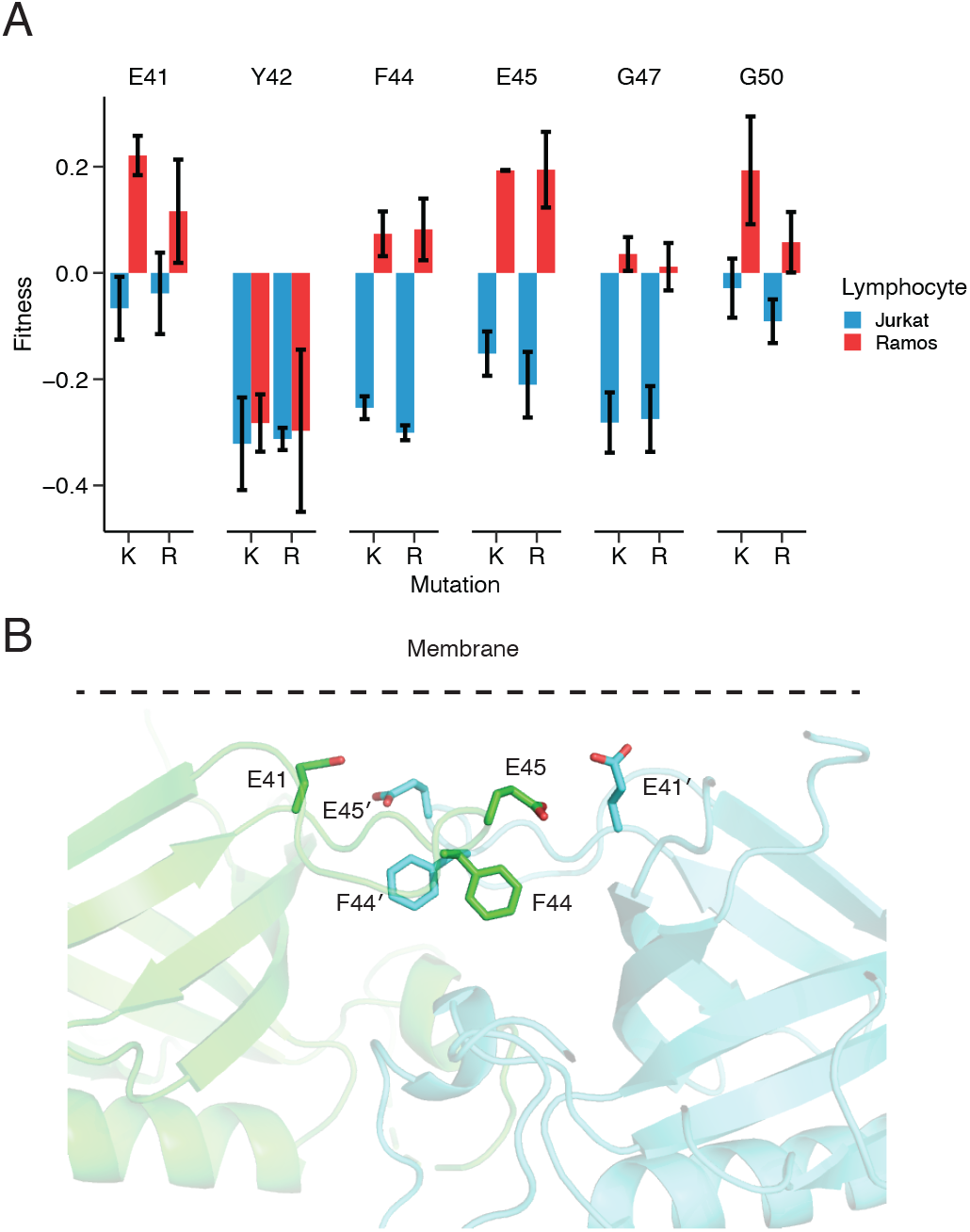
Substitutions of positively charged residues near the plasma membrane have differential fitness effects. (A) A set of mutations that show differential fitness in Ramos and Jurkat cells. Fitness scores in Jurkat cells (blue) are shown alongside fitness scores in Ramos cells (red) for lysine and arginine substitutions at select positions in the S1 loop. (B) Positioning of the residues that exhibit differential fitness. Stick representation of residues that change their fitness in the two cell lines (PDB: 1BTK). Note that Gly^47^ and Gly^50^ are not shown because they have no side chains. The predicted positioning of the plasma membrane is indicated by a dashed line.

Instead, mutations at these sites are almost always loss of function in both the Ramos and Jurkat cells, consistent with these residues being critical for membrane binding (Figure S6).

The differential fitness in the two contexts suggests that in Ramos cells, but not in Jurkat cells, BTK activity is enhanced by increasing membrane localization through increased positive charge (or decreased negative charge) in the PH domain. This idea is supported by the observation that the residues that enhance fitness in the Ramos context when mutated to lysine or arginine are predicted to face toward the plasma membrane, based on structures of the BTK PH– TH module (Figure 7B). One such residue is Glu^41^, for which the mutation E41K is activating in Ramos cells [6]. E45K, like E41K is also strongly activating in Ramos cells. These substitutions have a very different effect in Jurkat cells. The E41K mutation results in a loss of function, and the E45K mutation results in an even greater loss-of-function in Jurkat cells. For this reason, we chose to examine E45K further.

### The mutation E45K in the BTK PH–TH module promotes membrane binding

To investigate the effect of mutations in the S1 loop on membrane binding, we measured the adsorption of the PH–TH module to membranes using purified components. We prepared supported phospholipid bilayers on glass coverslips with 96% 1,2-dioleoyl-sn-glycero-3-phosphocholine (DOPC) and 4% PIP_3_. As a control, we used bilayers containing 4% phosphatidylinositol (4,5)-bisphosphate (PIP_2_) instead of PIP_3_. We purified three variants of the BTK PH–TH module (residues 1–171): PH–TH^WT^, PH–TH^E45K^ (the activating variant discussed above), and PH–TH^Y42K^ (the mutation at the dimer interface that we suspect destabilizes the PH domain), all tagged with mNeon-Green at the C terminus. Bilayer adsorption was quantified by imaging using total internal reflection fluorescence (TIRF) microscopy, which can discriminate between protein in solution and on the bilayer, as described previously for the BTK PH–TH module [62] (Figure 8A).

When using low solution concentrations of the PH–TH module (5 pM), and imaging using continuous monitoring and a 20 ms exposure, we observed single PH–TH^WT^ molecules diffusing freely across the bilayer. Membrane-bound dimers are not formed at this concentration. Quantifying the numbers of proteins adsorbed after a 10-min pre-incubation showed that PH– TH^E45K^ adsorbs significantly more quickly than PH–TH^WT^ to PIP_3_ bilayers (the mean fold change in the number of spots compared to the wild-type protein is 1.7 ± 0.24, Figure 8B). PH– TH^Y42K^ adsorbs significantly less quickly than PH–TH^WT^ to PIP_3_ bilayers (mean fold change of 0.23 ± 0.23). The different membrane adsorption rates between these three proteins are not due to differences in photobleaching (Figure S8A). All three proteins have similar diffusion rates, as measured by per-frame displacements, indicating that different oligomeric states do not confound the differences in adsorption at the low surface densities used in this experiment (Figure S8B).

**Figure 8.**
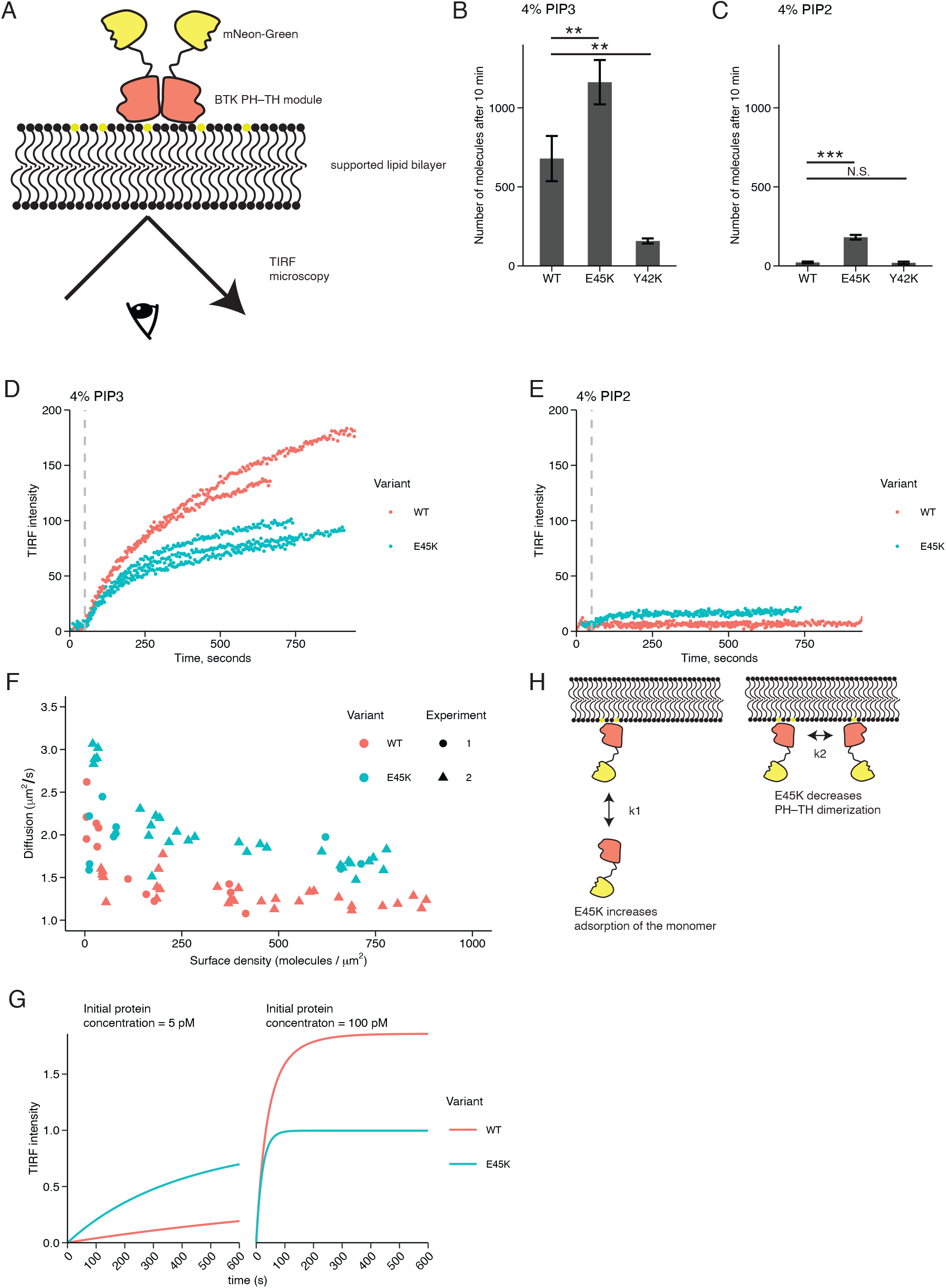
The mutation E45K increases the ability of monomeric BTK PH–TH module to bind membranes but decreases PH–TH dimerization. (A) Schematic of the adsorption assay. Purified PH–TH modules with a C-terminal EGFP tag adsorbed onto supported-lipid bilayers containing either 4% PIP_3_ or 4% PIP_2_ and 96% DOPC. Adsorption was visualized with TIRF microscopy which excites only the membrane-bound molecules. (B) PH–TH^E45K^ at low concentrations adsorbs more than PH‒TH^WT^ to bilayers containing 4% PIP_3_. The mean ± standard deviation number of molecules adsorbed after incubating 5 pM of protein (PH‒TH^WT^, PH‒TH^E45K^, PH‒TH^Y42K^) for 10 min was measured using trackmate [75] in Fiji. N = 4 trials. N.S: not significant, *p < 0.05, **p < 0.01, ***p < 0.001, t test. (C) PH–TH^E45K^ adsorbs more than PH‒TH^WT^ to bilayers containing 4% PIP_2_. Otherwise as in (B). (D) PH–TH^E45K^ adsorbs less than PH‒TH^WT^ to bilayers containing 4% PIP_3_ at high solution concentrations. Average TIRF intensity across the imaging field was measured after injection of 100 pM PH–TH^E45K^ or PH‒TH^WT^ (key). The vertical dashed line represents the time at which the protein was injected. The different curves represent independent trials (n = 2 for wild-type, n = 3 for E45K). (E) PH–TH^E45K^ at high concentrations adsorbs more than PH‒ TH^WT^ to bilayers containing 4% PIP_2_. Otherwise as in (D). (F) PH–TH^E45K^ exhibits faster diffusion than PH‒TH^WT^ at high surface densities. The diffusion rate (y axis) was measured for different protein densities on 4% PIP_3_-containing bilayers (x axis) using fluorescence correlation spectroscopy. Measurements were performed for PH–TH^WT^ and PH–TH^E45K^ in two independent experiments (key). (G) Kinetic modeling of PH–TH^E45K^ and PH–TH^WT^ proteins (key) at two different concentrations. The monomeric association (*k*_1_) rate constant was 4.7 x 10^-4^ pmol^-1^sec^-1^ for both mutants [5] and the dimer dissociation rate constant (*k*_–2_) was 5.0 x 10^-4^ sec^-1^ for both mutants. The monomeric disassociation (*k*_–1_) rate constants were 1.5 and 1.5 x 10^-4^ sec^-1^ for PH– TH^WT^ and PH–TH^E45K^, respectively. The dimeric association rate constant (*k*_2_) was 100 pmol^-1^sec^-1^ and 0 pmol^-1^sec^-1^ for PH–TH^WT^ and PH–TH^E45K^, respectively. (H) A model for how the mutation E45K changes the rate constants associated with monomer adsorption and dimerization.

As noted earlier, the Y42K mutation had been implicated in PH–TH dimerization, but these results show that the mutation most likely affects structural stability. At these concentrations, dimerization is not relevant, so it seems likely that the population of folded PH–TH^Y42K^ proteins is decreased. PH–TH^E45K^ and PH–TH^WT^ have the same distribution of intensities for all measured particles, arguing against differences in the ability to find and count the particles (Figure S8C). The PH–TH^Y42K^ protein shows a slight decrease in particle intensity, perhaps because of effects on the orientation or distance between the fluorescent protein and PH–TH module.

The PH–TH module of BTK does not bind with high affinity to membranes that contain PIP_2_ instead of PIP_3_ [5] (Figure 8C). Strikingly, the PH–TH^E45K^ mutant has significant adsorption rates to PIP_2_ bilayers (mean fold change compared to PH–TH^WT^ = 8.2 ± 0.27). Loss of lipid specificity has been reported for PH–TH^E41K^ [6], and Glu^41^ is located close to Glu^45^. These data provide a mechanism for the increased fitness that we observe in the Ramos system, because PH–TH^E45K^ adsorbs more than PH–TH^WT^ to membranes containing either PIP_2_ or PIP_3_.

Glu^45^ is not conserved across the PH–TH modules of the human Tec kinases but is only present in BTK (Figure S8D). This suggests that Glu^45^ was selected in the BTK lineage to increase the specificity of the BTK PH–TH module for PIP_3_ even at the cost of a loss in affinity for the plasma membrane. Indeed, substitution of Glu^45^ to almost any other residue (not just lysine or arginine) enhances BTK fitness in Ramos cells (Figure S6A), perhaps by alleviating the membrane repulsion induced by the negatively charged glutamate. ITK does not have a glutamate at this position and in some fish species (e.g. *Esox lucius* or *Oryzias latipes*) even has an arginine at this position. ITK is more constitutively membrane-localized than BTK [34], suggesting that the PH–TH module of ITK does not require the same restraint on its membrane localization.

### The mutation E45K disrupts dimerization of the BTK PH–TH module

The increased membrane binding rates of PH–TH^E45K^ compared to PH–TH^WT^ provides an explanation for how the mutation E45K increases fitness in Ramos cells but this fails to explain how the same mutation decreases fitness in Jurkat cells. We wondered whether the mutation E45K decreases fitness in Jurkat cells by decreasing PH–TH dimerization because this residue is located close to the membrane interface. To examine this, we increased the solution concentration of the PH–TH proteins in our reconstitution experiment to promote higher surface densities and dimerization. In the cell, the membrane-bound concentration of BTK is modulated by PIP_3_ levels, but changing the PH–TH solution concentration is a convenient way to change the membrane-bound concentration in our reconstitution system.

We used TIRF intensity to measure PH–TH adsorption at higher concentrations rather than using single-molecule counting, which is not possible at higher concentrations. At higher protein solution concentrations (100 pM, Figure 8D or 250 pM, Figure S8E), PH–TH^WT^ adsorbs more than PH–TH^E45K^ to bilayers containing PIP_3_. This is in striking contrast to the results obtained when solution concentration of the PH–TH module are low (Figure 8B). These results are consistent with decreased PH–TH dimerization in the E45K mutant. Importantly, even at high solution concentrations, PH–TH^WT^ fails to adsorb to bilayers containing PIP_2_ (Figure 8E).

We turned to fluorescence correlation spectroscopy (FCS) to examine further the effect of the E45K mutation on PH–TH dimerization. FCS measures fluctuations in the fluorescence intensity as molecules diffuse in and out of a small illumination volume. These data can be used to produce an autocorrelation function, from which surface densities and diffusion rates of the fluorescent molecules can be determined. FCS analyses have been used previously to show that PH–TH^WT^ dimerizes [5] by using the fact that molecular dimers on the membrane surface exhibit notably slower diffusion than monomers. By measuring the mean surface diffusion rate of membrane bound proteins as a function of surface density, it is possible to map a two-dimensional monomer-dimer binding equilibrium on the membrane surface [63]. Consistent with this, the two-dimensional diffusion rate of PH–TH^WT^ adsorbed onto PIP_3_-containing bilayers is lower at high surface densities (Figure 8F). By contrast, PH–TH^E45K^ diffuses rapidly at high surface densities (Figure 8F). At low surface densities, significantly below the two-dimensional dimerization dissociation equilibrium constant, both proteins show similar and rapid diffusion, which is also consistent with the lack of difference in diffusion rates observed using single-molecule TIRF (Figure S8C). These FCS data are consistent with the mutation E45K decreasing PH–TH dimerization.

A kinetic model captures how a mutation that disrupts dimerization might have different effects depending on the degree of dimerization in the system. We used a two-step model in which the PH–TH protein first binds PIP_3_ and then the PH–TH::PIP_3_ complex dimerizes (Figure S8F). To model dimerization of PH–TH^E45K^, we decreased the rate constant for PIP_3_ dissociation form monomer PH–TH^E45K^ (*k*_–1_) and decreased the association rate constant of the two protomers of the dimer (*k*_2_). These adjustments to the rate constants result in increased binding of PH– TH^E45K^ to PIP_3_, relative to PH–TH^WT^, when the system is initialized with low protein concentrations (e.g. 5 pM). When the system is initialized with high protein concentrations (e.g. 100 pM), the same adjustments decrease the binding of PH–TH^E45K^ to PIP_3_, relative to PH–TH^WT^(Figure 8G). These results explain both the loss-of-function due to the E45K mutation in Jurkat cells (which depend on dimerization) and the gain-of-function caused by the same mutation in Ramos cells (which do not require PH–TH dimerization) (Figure 8H).

## Discussion

In this study, we leverage the power of high-throughput mutagenesis to examine the importance of PH‒TH dimerization and kinase activity for BTK function in lymphocytes. We develop mutagenesis assays in two cell lines: Ramos B cells and Jurkat T cells. In Jurkat cells, we find that PH–TH dimerization is required for signaling, with a landscape of mutations in the PH–TH dimer interface that is consistent with predictions from the crystal structure of the Saraste dimer [9]. We also find that mutations of key catalytic residues in the kinase domain abolish CD69 expression, indicating that BTK signaling requires its kinase activity. In contrast, we find no evidence for the importance of PH–TH dimerization or kinase activity in Ramos cells. Instead, substitution of neutral and negatively charged residues by lysine and arginine at positions that are predicted to interact with the plasma membrane increase BTK fitness. In Ramos cells, BTK signaling does not depend on dimerization of the PH–TH module, whereas this dimerization is critical in Jurkat cells. These differences are underscored by the E45K mutation in the PH domain, which increases fitness in Ramos cells but decreases fitness in Jurkat cells. We find that this mutation enhances adsorption of the PH–TH module on membranes while decreasing PH– TH dimerization.

Collectively, the data presented here show that BTK signals in two modes. In one mode, as seen for Jurkat cells, BTK kinase activity and PH–TH dimerization are both important. In another mode, as seen for Ramos cells, kinase activity and PH–TH dimerization are both dispensable. But Ramos cells still need BTK for signaling, and our work shows that membrane binding by the PH domain is important, which suggests that BTK plays a scaffolding role in this context.

The catalytic function of tyrosine kinases have gained the most attention, because kinase activity is commonly dysregulated in cancers and readily blocked by kinase inhibitors. Even though the kinase-independent signaling functions of protein kinases were discovered early in the study of oncogene function [67], these functions have received relatively little attention. It is evident from the modular domain structure of cytoplasmic tyrosine kinases that they have a powerful capacity to form networks of interactions that can control cell signaling through colocalization and liquid–liquid phase separation. The recent realization that BTK acquires drug-resistance mutations that disrupt its kinase activity refocuses our attention on the scaffolding functions of BTK and other cytoplasmic tyrosine kinases. Our work shows that mechanisms that have evolved to support kinase activity, such as the PH–TH dimerization that is a feature of BTK, may be separable from those that evolved to support scaffold activity. Future drug development efforts might benefit from a search for molecules that can tease apart the many signaling modes of kinases.

Our phylogenetic analysis of Tec-family kinase sequences shows that the structural elements involved in PH–TH dimerization, which are by extension also important for kinase activity, were not present in the first Tec kinases or BTK proteins. Instead, the signature for PH– TH dimerization appears later, after the divergence of the ray-finned fishes from other jawed vertebrates. What evolutionary pressure might have led to PH–TH dimerization in BTK? Comparing BTK from zebrafish (*Danio rerio*, of the class Actinopterygii, which lacks the PH– TH dimerization signature) and the frog (*Xenopus laevis*, of the class Amphibia, which has the PH–TH dimerization signature) offers clues. In the zebrafish, there are no clearly defined regions of B and T-cell association that form in response to infection [64]. By contrast, in the frog, B-cell follicles form in the spleen and are surrounded by a peripheral layer of T-cells in a structure resembling the germinal center [65]. Both the B and T cells in this structure are presented antigen by a primordial follicular dendritic cell [65]. Thus, amphibians mark the dawn of a new type of antigen presentation to B cells. We imagine that this new format of antigen presentation required changes in the timing and location of B-cell activation, perhaps selecting for BTK PH–TH dimerization as a new level of control in the process. Indeed, in mammals, the format of antigen presentation (whether on cells, in solution, or on viruses) determines the timing and degree of PIP_3_ accumulation and the nature of B-cell signaling [66].

### Methods Cell lines

Jurkat T cells and ITK-deficient Jurkat cells (a kind gift from Wan-Lin Lo) were cultured in 5% fetal bovine serum (FBS) in RPMI media without antibiotics supplemented with 1x glutamax (Thermo Fisher). Ramos RA1 B cells (ATCC) and BTK-deficient Ramos cells were cultured in 10% FBS in RPMI supplemented with 1x glutamax. HEK293FT cells (Thermo Fisher) were cultured in 10% FBS in DMEM supplemented with 1x glutamax. All cells were maintained in a humidified incubator at 37°C and 5% CO_2_.

### Generation of BTK-deficient Ramos B cells

BTK-knockout Ramos B cells were generated by Crispr/Cas9 technology. sgRNAs targeting BTK (ordered as a pair with its reverse complement, oligos #336 and #337, Table S1) or a non-targeting control (oligos #340 and #341, Table S1) were cloned into the pU6-(BbsI)_CBh-Cas9-T2A-BFP vector (Addgene Plasmid #64323) as previously described [68]. 40 µg of plasmid was then electroporated into 10 million Ramos RA1 cells using an electroporator (BioRad Gene Pulser Xcell) with a 0.4 cm cuvette and the following settings: 400 µF, 250 V, 1000 Ω using an exponential-decay pulse. Cells were allowed to recover for three days and then sorted to single cells into a 96-well plate based on blue fluorescent protein (BFP) fluorescence using a Sony SH800 cell sorter and a 100 µm sort chip in culture media supplemented with penicillin/streptomcyin. Clones were grown for ∼2 weeks or until the media turned orange, then 200 µL of media was added and cells were grown for an additional week. Clones were then plated up to a 24-well plate in replicate plates, and one of the replicates was used for western-blot analysis. Clones that were BTK-negative were expanded and frozen, along with some non-targeting controls.

### Lentiviral production and titering

To prepare lentivirus, HEK293FT cells were seeded at 250,000 cells/mL in 5 mL in 6-cm dishes in culture media (day 1). The next day (day 2), cells were transfected with a mixture of transfer plasmid (modified from pHIV-EGFP, addgene #21373, 5 µg) and two packaging plasmids (pMDG.2, #12259, 1.25 µg and psPAX2, #12260, 3.75 µg) in 500 µL of opti-mem (Thermo Fisher) using lipofectamine LTX (Thermo Fisher) according to the manufacturer’s instructions. The following morning (day 3) cells were re-fed. The following day (day 4) 5 mL of viral supernatant was harvested and stored at 4°C, and the cells were re-fed. The following day (day 5) an additional 5 mL of viral supernatant was harvested and combined with the previous day’s aliquot for a total of 10 mL of viral supernatant. This was then concentrated using Lenti-X concentrator (Takara Bio) according to the manufacturer’s instructions, and the pellet from this concentration was resuspended in 1 mL of RPMI + 5% FBS, aliquoted, flash frozen in liquid nitrogen, and stored at –80°C.

Lentiviruses were titered by thawing an aliquot and adding 0, 5, 10, 20, 40, or 80 µL of virus to 500 µL ITK-deficient Jurkat T cells in a 24 well plate, along with 10 µg/mL polybrene. The following day, cells were centrifuged (300g for 5 min) and resuspended in fresh media, then diluted 2x and plated onto a 96-well plate. Cells were grown for two additional days and then the plate was analyzed for GFP fluorescence using an Attune flow cytometer (Thermo Fisher) equipped with a 96-well plate autosampler. The fraction of GFP positive cells as a function of concentration was fit to the standard binding isotherm to obtain titer values.

### Library cloning

Saturation-mutagenesis libraries corresponding to regions of interest in BTK were cloned as follows. An oligo pool targeting the region of interest and containing the human wild-type sequence, additional wild-type sequences with random synonymous-codon substitutions, two codon substitutions for each amino acid (except for methionine and tryptophan), and two stop codons at each position was designed and purchased from Twist Biosciences. This oligo pool was designed to have BsaI cloning sites flanking the insert, and sequences were re-coded using synonymous substitution if they contained BsaI, XmaI, or BamHI sites.

Upon receipt, the pool was amplified using PCR for 15-20 cycles in a 50 µL reaction. The two flanking constant regions of BTK (the left and right sides of the insert, Table S1) were also amplified by PCR using primers that added insert-compatible BsaI sites. The three fragments were then column-purified (Zymo DNA Clean and Concentrator) and ligated in a combined digestion-ligation step first at 37°C for 2 h and then at 16°C overnight with 1 µL BsaI-HF, 1 µL T4 DNA ligase, 1x T4 DNA ligation buffer, and 20% polyethylene glycol (New England Biolabs) in a total reaction volume of 50 µL. The next day, the reaction was heat-inactivated at 65°C for 10 min followed by column purification. The sample was then digested for 1 h at 37°C using XmaI and BamHI in 1x cutsmart buffer (New England Biolabs). The plasmid backbone (pHIV-EGFP addgene #21373) was digested in a separate 50 µL reaction. The sample insert and backbone were then gel purified on a 1% agarose gel.

Following gel extraction and purification using a gel-extraction kit (Qiagen), insert and vector were ligated for 1 h at room temperature in a 20 µL reaction containing 1x T4 DNA ligase buffer and 1 µL T4 DNA ligase, along with a no-insert control. Ligation reactions were column-purified (Zymo DNA Clean and Concentrator) and eluted in 6 µL. 3 µL of this reaction was used to electroporate Endura electrocompetent cells (Biosearch technologies) according to the manufacturer’s instructions, slightly modified as follows. Immediately after electroporation, cells were resuspended in 1 mL of recovery buffer (Biosearch technologies) and grown for 1 h at 30°C while shaking. This volume was then diluted to 50 mL, and 50 µL were plated onto LB-agar plates containing ampicillin (100 µg/mL). The remaining volume was grown overnight at 30°C.

The next day, the plates were examined. The experiment was only continued if the library coverage was at least 500x (for example, a 500-variant library needs 250,000 total transformants, or 250 colonies on the sample plate, with little background on the control plate). If the transformation was successful, plasmid DNA was extracted from the 50 mL cultures using a midiprep kit (Qiagen) according to the manufacturer’s instructions, and used to prepare lentivirus.

### Cell analysis and selection

To measure BTK-dependent CD69 expression using BTK-variant libraries, ITK-deficient Jurkat T cells or BTK-deficient Ramos B cells were plated in 10 mL at a density of 500,000 cells/mL in 10 cm dishes in triplicates or quadruplicates. The cells were then transduced on day 1 with titered lentiviral libraries, adding a volume of virus such that at most 25% of the cells were GFP positive (corresponding to < 3% of cells having a multiplicity of infection > 1) and 10 µg/mL polybrene.

The following morning (day 2) cells were re-fed by centrifugation at 300g for 5 min followed by resuspension in fresh media, with a 2x dilution, and re-plated. Cells were allowed to recover for one additional day (day 3). On day 4, Ramos cells were stimulated in the evening with 4 µg/mL anti-IgM (I2386, Sigma-Aldrich).

On day 5, Ramos or Jurkat cells were stained with anti-CD69 as follows. Cells were concentrated by centrifugation at 300g for 5 min, then resuspended in 200 µL of cell-staining buffer (phosphate-buffered saline containing 10% FBS and 0.05% w/v sodium azide) containing a 1:20 dilution of PerCP-Cy5.5 conjugated anti-CD69 (Cell Signaling Technology #28633). Cells were then incubated on ice for 30 min in the dark. Following this incubation, cells were centrifuged again and washed with 500 µL cell-staining buffer, then centrifuged and resuspended in 1 mL of cell-staining buffer for sorting. After this staining protocol, 100 µL (10%) of sample was removed as input and kept on ice during sorting. Cells were sorted for a GFP-positive and CD69-positive population on a Sony SH800 Cell Sorter, gating based on single and double negative controls, sorting at least 200x the number of cells as variants in the library.

Following sorting, input and sorted samples were mixed with 1 mL of TRI reagent (Sigma-Aldrich), concentrating the sorted sample by centrifugation (300g, 5 min) if the volume exceeded 200 µL. RNA was extracted according to the manufacturer’s protocol, except that after the aqueous phase was transferred to a new tube, it was mixed 1:1 with chloroform and centrifuged at 21,000g for 10 min at 4°C as an additional wash step. 4 µL of linear acrylamide (Thermo Fisher) was added as carrier.

For some experiments, cell sorting was not required but the fractions of CD69 and GFP positive cells were still analyzed. For these experiments, the Attune flow cytometer was used instead of the Sony SH800, and the protocol for staining cells was exactly above, with one modification. Because of the different laser and filter configurations on the sort instruments, the staining antibody used for analysis was APC-conjugated anti-CD69 from Biolegend, #310910, which is the same clone as the other conjugate (Cell Signaling Technology #28633).

In experiments using the HRD-motif libraries, CD69-positive cells were separated with a MACS kit instead of a cell sorter, using the CD69 microbread kit II for human (Miltenyi Biotec), according to the manufacturer’s instructions.

### Library preparation and sequencing

RNA-seq libraries were prepared from TRI reagent-extracted samples as follows, beginning with the reverse-transcription. Following precipitation, RNA was resuspended in water containing 5 µM RT primer (oligo #296, Table S1). The mixture was heated to 65°C for 5 min, then snap cooled on ice. The RNA–oligo mixture was then mixed with 1x first-strand buffer (Thermo Fisher), 0.5 mM deoxynucleotide triphosphates, 10 mM DTT, 1 µL SuperaseIn (Thermo Fisher), and 0.5 µL Superscript III (Thermo Fisher). The final reaction mixture was incubated at 50°C for 1 h. Followed this incubation, RNA was hydrolyzed by addition of 5 µL of 1 M NaOH and incubation at 90°C for 10 min. The mixture was then neutralized with 25 µL of 1 M HEPES, pH 7.4, and desalted using a Micro Bio-Spin P-30 column, in tris (BioRad), eluting the cDNA in 60 µL.

The cDNA was PCR amplified in one round (the S1 loop, S2 loop, and activation loop libraries) or two rounds (the HRD-motif library and the peripheral and canonical-site libraries). In the one-round protocol, long primers that were specific to the template but also added the Illumina adapters, flow-cell hybridizing sequences, and barcodes were used (oligos #209, #210, #211, #212, #213, #214, #215, #216, #217, #218, #219, #220, #221, #222, #223, #224, and #225, Table S1). In the two-round protocol, the first-round PCR added the priming regions for Illumina sequencing but did not add the barcodes (using primers #323, #324, #328, #329, #391, and #392, Table S1). The second-round PCR used primers that annealed to the Illumina priming regions, adding the flow-cell hybridizing sequences and barcodes (using primers BC_p5_1, BC_p5_2, BC_p5_3, BC_p5_4, BC_p5_5, BC_p5_6, BC_p5_7, BC_p5_8, BC_p7_1, BC_p7_2, BC_p7_3, BC_p7_4, BC_p7_5, BC_p7_6, BC_p7_7, BC_p7_8, BC_p7_9, BC_p7_10, BC_p7_11, and BC_p7_12, Table S1). Amplified libraries were gel-purified on a 2% agarose gel, extracted with a gel-extraction kit (Qiagen), pooled at equal molar based on the band intensities, and then concentrated using a clean and concentrator kit (Zymo). Libraries were sequenced using a MiSeq sequencer and V2 chemistry with paired-end 150 by 150 base reads.

### Data analysis

Fastq files from MiSeq runs were aligned to the Fasta files containing the full sequences of each variant using Kallisto [69] to generate read counts for each variant. The next steps of data processing proceeded independently for each pair of input and sorted datasets. To generate fitness scores, a read cutoff of 50 reads was first applied to the input libraries such that any variant not passing this threshold was discarded. Next, the unnormalized scores were calculated by dividing the number of reads in the sorted dataset by the number of reads in the input dataset and taking the log_10_. These unnormalized scores were normalized by subtracting the mean of the wild-type fitness scores, according the equation 1:

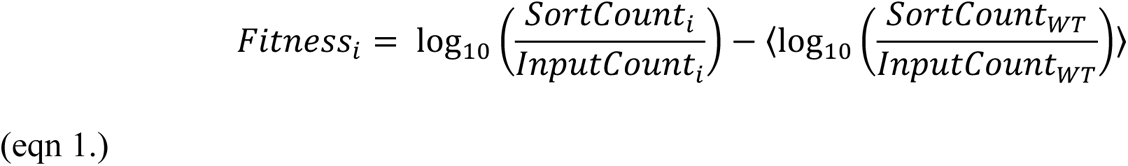

In equation 1, *Fitness_i_* denotes the fitness score for a particular variant *i*.

The number of wild-type sequences was 20–30, based on the number of positions in the library: because two codons were included for each amino acid that has at least two codons, the number of wild-type variants was the sum of the number of positions and additional sequences with multiple random synonymous substitutions. The fitness scores are plotted ± standard error.

Code to generate saturation-mutagenesis sequences and to analyze RNA-seq libraries was written using R [70] and Python and is available on Github (https://github.com/timeisen/MutagenesisPlotCode).

### Bacterial Expression and Purification of PH–TH modules

The BTK PH–TH module and variants were cloned by ordering a version of the BTK coding sequence (residues 1–171) codon optimized for *E. coli* using the codon-optimization tool (Integrated DNA Technologies). This sequence was PCR amplified (oligos #246 and #421, Table S1) and cloned into the pET-SUMO vector, as described previously [4]. The E45K and Y45K mutations were introduced into the PH–TH module by overlap-extension PCR (oligos #414 and #420, Tabls S1). The final construct contained an N-terminal hexahistidine tag followed by a SUMO tag, then the PH–TH module, and a C-terminal mNeonGreen, under the control of a T7 promoter.

The final plasmid was transformed into BL21-DE3 *E. coli* and plated onto kanamycin-resistant plates. Colonies were used to start a 50 mL liquid culture which was grown overnight at 37°C in terrific broth (TB) containing 50 µg/mL kanamycin. The following morning, 1 mL of the saturated culture was used to start a 1 L culture (TB and 50 µg/mL kanamycin), which was grown until it reached an OD_600_ of 1.5. At this point, protein production was induced by diluting the culture 2-fold with TB containing 50 µg/mL kanamycin, 2 mM Isopropyl ß-D-1-thiogalactopyranoside (IPTG) and 200 µM ZnCl_2_, pre-cooled to 4°C. The 2 L culture was then grown overnight at 18°C.

The next morning, the bacterial were pelleted by centrifuging at 4000 rpm for 15 min at 4°C in a swinging-bucket centrifuge. The pellet was resuspended in 10 mL of lysis buffer (20 mM tris, pH 8.5, 500 mM NaCl, 20 mM imidazole, and 5% glycerol supplemented with tablet protease-inhibitors [Roche]). The bacteria were lysed by sonication and the supernatant was cleared by centrifugation at 16,500 rpm for 30 min at 4°C.

The clarified supernatant was loaded onto a HisTrap FF column (Cytiva) using a peristaltic pump and washed with 10 column volumes of lysis buffer. The loaded column was then transferred to an ÄKTA System FPLC (Cytiva) and washed with an additional 10 column volumes of lysis buffer. Protein was eluted in a single step using elution buffer (the lysis buffer supplemented with 400 mM imidazole). Following elution, the sample was concentrated down to 0.5 mL using a 30 kDa concentrator (Amicon) and loaded onto a HiTrap desalting column (Cytiva), eluting with gel-filtration buffer (25 mM tris, pH 8.0, 150 mM NaCl, 1 mM tris(2-carboxyethyl)phosphine and 5% glycerol). The sample was then diluted to 20 mL, combined with 25 units of SUMO protease (Sigma-Aldrich), and incubated overnight at 4°C.

Cleaved protein was loaded onto a HisTrap FF column (Cytiva). The protein was eluted with lysis buffer: although the protein no longer had the hexahistidine tag, PH domains have slight affinity for the Ni-NTA column and do not elute with the flow-through. The eluted protein was concentrated using a 10 kDa concentrator (Cytiva) down to 0.5 mL and loaded onto an Superdex 75 Increase 10/300 GL (Cytiva). The major peak, eluting at ∼56 mL was collected and concentrated using a 10 kDa concentrator to ∼30 µM, flash frozen in liquid nitrogen, and stored. Typical yields were 1–2 nmol.

### Supported-lipid bilayer preparation

Supported-lipid bilayers were prepared essentially as described [62] with slight modification. Lipids were prepared by mixing 96% 18:1 1,2-dioleoyl-sin-glycero-3-phosphocholine (DOPC) (Avanti Polar Lipids) and either 4% PIP_3_ (Echelon Biosciences, Inc) or 4% PIP_2_ (Brain PI(4,5)P2, Avanti) in an etched 50 mL round-bottom flask. To assess lipid quality, 0.005% TRITC-DHPE (Biotium) was added. PIP_3_ was dissolved in a mixture of 1:2:0.8 chloroform to methanol to water just prior to use.

The solvent from the mixture was evaporated using rotary evaporation and then further dried under nitrogen for 15 min. Dried lipids were stored overnight at 4°C. The next morning, lipids were rehydrated by dissolving to 1 mg/mL in water by vortexing. Small unilamellar vesicles (SUVs) were prepared by sonication (Analis Ultrasonic Processor 750 watt, 20 kHz) at 33% power, 20 s on, 50 s off for a total of 1 min 40 s active sonication. SUVs were then diluted to 0.25 mg/mL in 0.5x tris buffered saline and assessed for uniformity and size using dynamic light scattering (Brookhaven Instrument Corporation).

80 µL of the SUV mixture was added to each well of an etched glass slide mounted onto a Sticky Slide VI 0.4 (Ibidi GMBH) and incubated for 30 min at room temperature. Each well was then washed with 1x hanks balanced salt solution (HBSS) under vacuum as described [62], then blocked for 10 min with β-Casein (0.1% w/v in HBSS) and washed again with 1x HBSS. Slides were then washed with imaging buffer (HBSS with 0.01% w/v β-Casein).

To image protein binding, 100 µL of imaging buffer were removed from the chamber (total volume 200 µL) and replaced with 100 µL of sample at the indicated times before acquisition. For single-molecule measurements and FCS, a scavenging mixture (10x: 100 µM β-mercaptoethanol, 20 mM Trolox, 3.2 mg/mL glucose oxidase [Serva Electrophoresis GMBH]) was prepared. This mixture was diluted 10-fold into the sample just before addition to the imaging chamber and imaging.

### TIRF imaging

TIRF microscopy was performed as described [5, 62]. Briefly, images were collected on a Nikon Eclipse Ti-inverted microscope (Nikon, Tokyo, Japan) with a 100 x TIRF objective (NA 1.49) and an Andor iXon electron-multiplying charge-coupled device (EMCCD) camera (Oxford Instruments). Lasers (488 and 561 nm; Coherent, Inc.) were used as illumination sources.

### Fluorescence correlation spectroscopy and analysis

FCS was performed as described [71]. Briefly, FCS was performed on a modified inverted microscope (Nikon TE2000), excited using bandpassed wavelengths from a white-light laser source (SuperK Extreme EXW-12, NKT Photonics, Copenhagen, Denmark). A 100x oil-immersion objective was used and signal was recorded with an avalanche photodiode detector (Hamamatsu). The signal was converted into autocorrelation signal by a digital correlator. Fitting to a 2D diffusion model was performed as previously described [5].

### Kinetic Modeling

Kinetic modeling was performed as previously described [72]. Briefly, a set of differential equations describing the reaction scheme were constructed in R and solved using deSolve [73]. Summary statistics and plotting were performed using ggplot2 [74].

### Western blotting

Western blotting was performed as described [62]. Blots were probed with a rabbit monoclonal antibody targeting BTK (Cell Signaling Technology #3533). For a loading control, blots were probed with a rabbit polyclonal antibody targeting 14-3-3 (Cell Signaling Technology #8312).

## Acknowledgments

We thank L. Nocka and S. Choi for helpful discussion, along with other members of the Kuriyan, Groves, and Weiss laboratories. We appreciate the technical advice of M. West, K. Pestal, H. Ren and W-L. Lo.

## Author Contributions

T.J.E, A.W., and J.K. conceived the project and designed the study. T.J.E. and S.G. performed the experiments. J.G. provided guidance and support for the microscopy experiments. T.J.E. wrote the paper, with review and editing from all authors.

## Data and materials availability

Raw fastq files and processed files are available on DRYAD. Code for generating saturation-mutagenesis libraries and analyzing them is available on Github (https://github.com/timeisen/MutagenesisPlotCode).

## Supplemental figure legends

**Figure S1.**
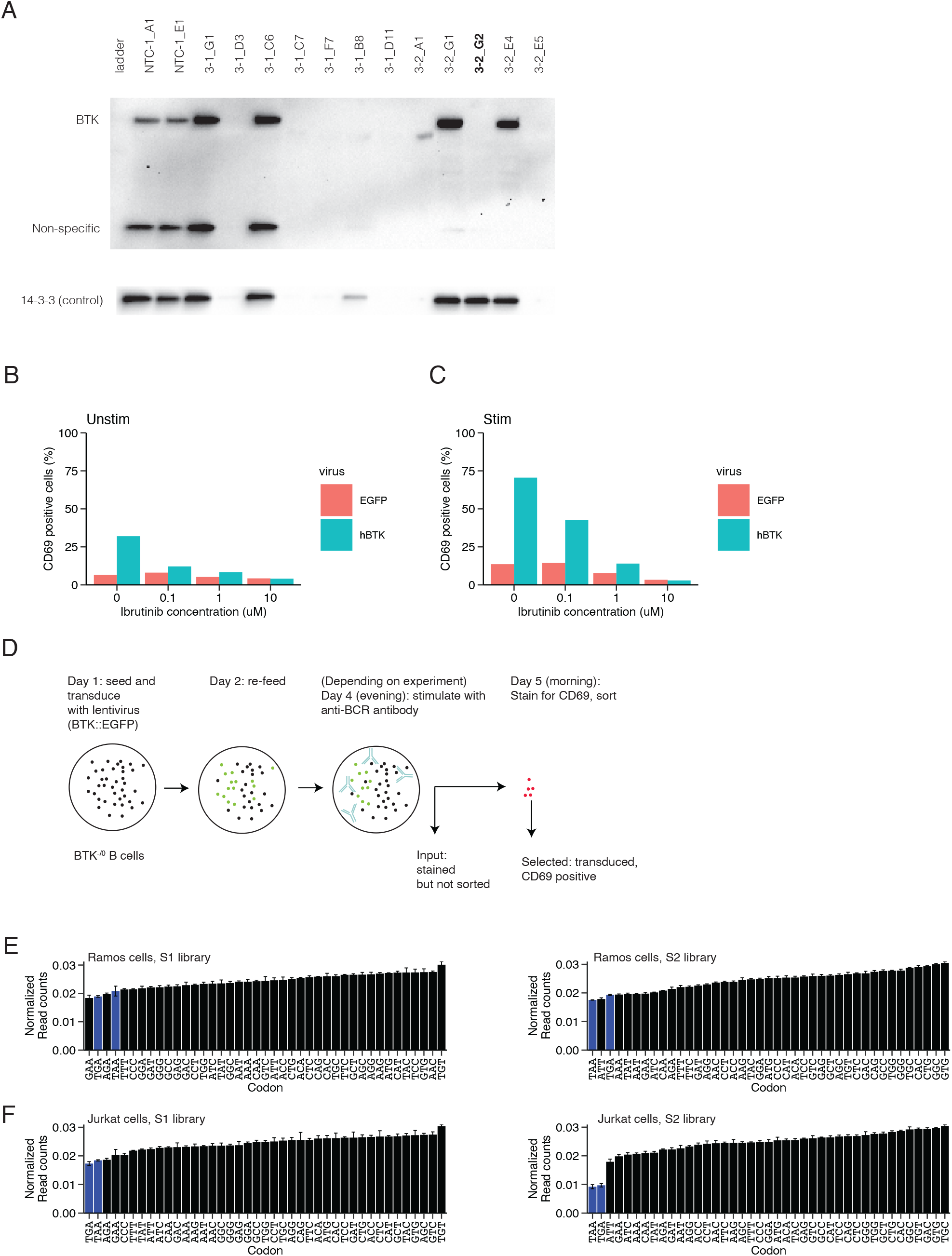
An assay for BTK fitness in Ramos B cells. (A) Western blot of BTK-knockout Ramos B-cell lines. The presence of BTK is indicated by the top band in the top blot for 14 CRISPR-targeted cell lines. Two of the lines used a non-targeting guide (NTC) and 12 of these used a guide targeting BTK exon 3. The cell numbers were not normalized prior to blotting: many of these lines failed to grow enough to produce signal, as judged by a loading control (anti-14-3-3, bottom blot). The BTK-deficient line 3-2-G2 was used in this study. (B) Ibrutinib decreases the BTK-dependent increase in CD69. ITK-deficient Jurkat cells were transduced with lentiviruses encoding human BTK or EGFP as in Figure 2C and then treated with ibrutinib at the indicated concentrations two days before sorting. Otherwise as in Figure 2C. (C) Ibrutinib decreases the stimulation and BTK-dependent CD69 expression. Otherwise as in (B). (D) Schematic showing the manipulations performed for screening Ramos cells for BTK function. BTK-deficient Ramos cells were transduced with viruses encoding wild-type BTK and mutations, and grown, and then sorted for CD69. (E) Stop codons are depleted in the input libraries in the Ramos-cell assay. The normalized read counts corresponding to codon substitutions at any position in the S1-loop library (left panel) or S2-loop library (right panel) in Ramos cells, shown as mean ± standard deviation for each of three replicates. The normalization is the raw read counts divided by the total number of reads in the library. The codons are ranked by the increasing order of read counts, and stop codons are shown in blue. (F) Stop codons are depleted in the input libraries in the Jurkat cell assay. Otherwise as in (E), with n = 4 biological replicates.

**Figure S2.**
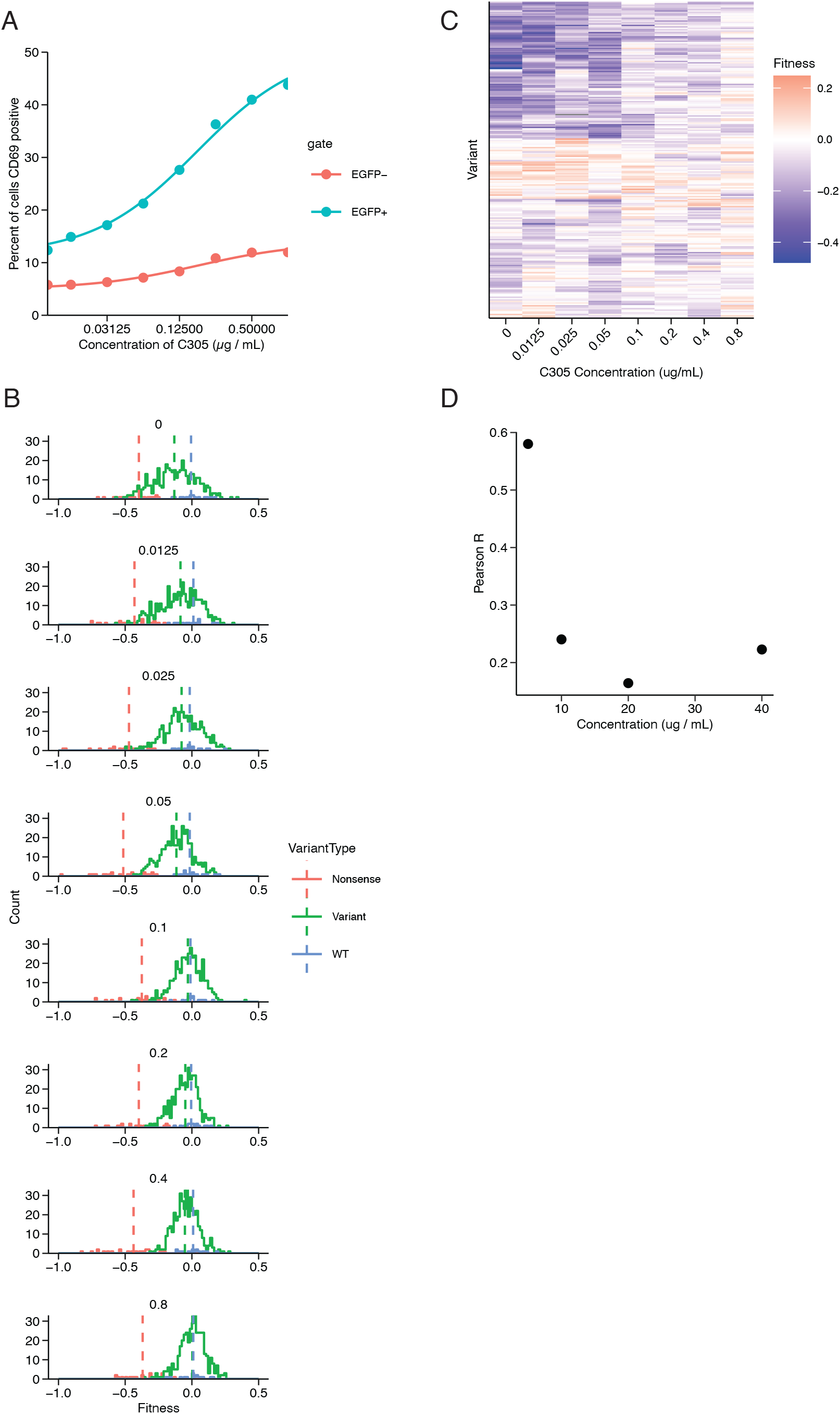
Stimulation promotes BTK-independent signaling in ITK-deficient Jurkat cells and BTK-deficient Ramos cells. (A) The fraction of CD69 positive ITK-deficient Jurkat cells as a function of increasing concentration of the anti-T cell receptor antibody C305, as measured by FACS. The percentages are shown for two sort gates: EGFP-positive cells are those that were transduced with the lentivirus expressing human BTK-IRES-EGFP, and EGFP-negative are untransduced cells. Points represent the measured values, and the line is a fit to a binding-isotherm model. (B) Fitness scores for all variants in the S1-loop library over increasing concentrations of C305. For each of the points in (A), mRNA-seq libraries were prepared from the input (unsorted) and CD69-sorted fractions of cells. The fitness values for the wild-type sequences (blue), sequences with a mutation (red), or sequences with a stop codon (key) are plotted as a histogram (key). The values above each histogram indicate the concentration of C305, in µg/mL. Over increasing concentrations, the nonsense variants resemble the wild-type variants but the stop-codon variants do not. (C) A heatmap showing the fitness values (key) for each variant in the S1-loop library over different concentrations of C305. Variants were clustered by Euclidian distance using the hclust algorithm in R. (D) Pearson correlation values between biological replicates for the S1-loop library in Ramos B cells over increasing concentrations of anti-IgM stimulatory antibody.

**Figure S3.**
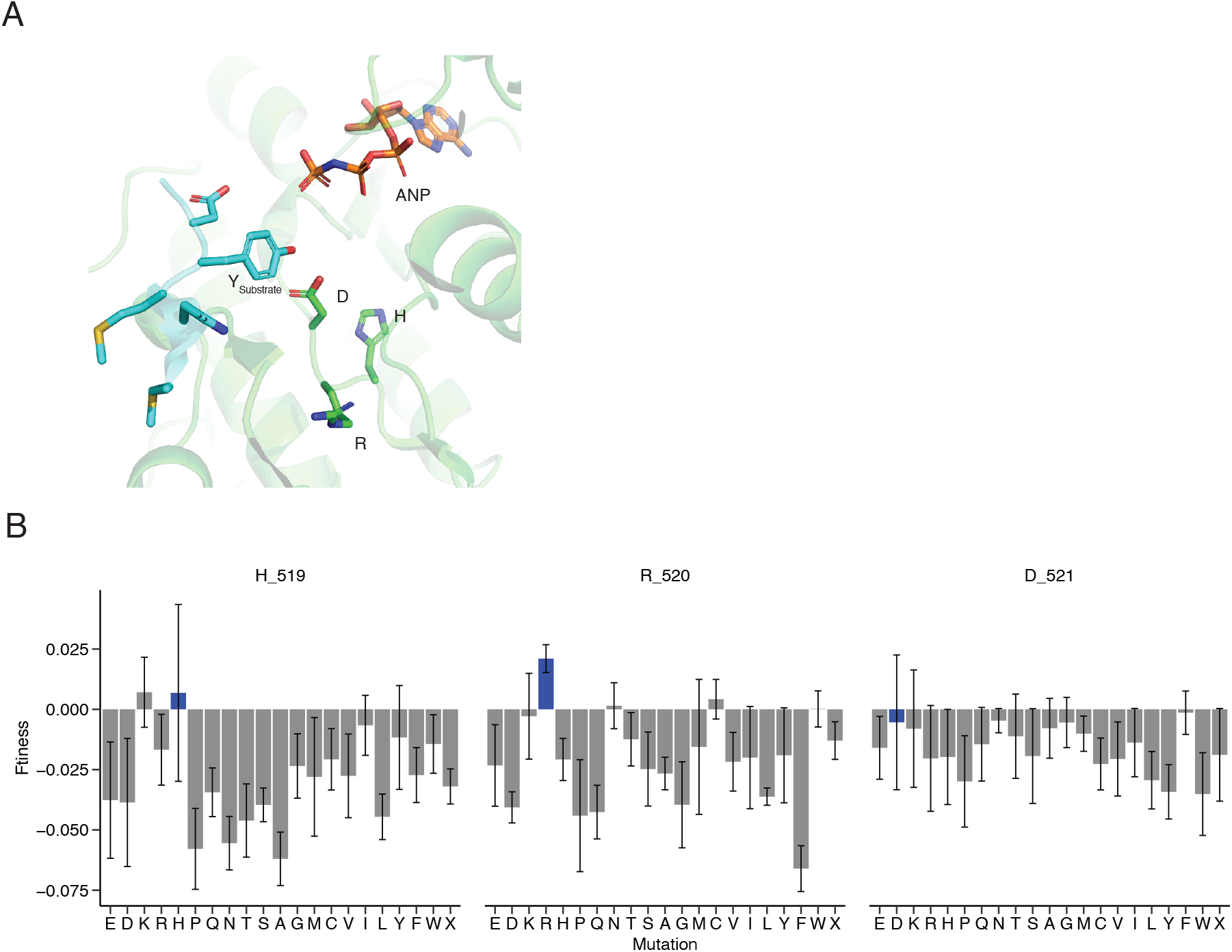
BTK kinase activity is dispensable in Ramos cells. (A) The HRD motif positions the catalytic base to abstract a proton from the peptide substrate. A crystal structure of the insulin receptor tyrosine kinase (PDB: 1IR3 [76]) in complex with phosphoaminophosphonic acid-adenylate ester. The histidine, arginine, and aspartate residues are shown as sticks (green) along with the ATP analog ANP (orange) and peptide substrate (cyan). (B) The HRD motif tolerates mutations. Fitness scores showing all possible residue substitutions to the HRD motif in Ramos cells. Otherwise as in Figure 2E.

**Figure S4.**
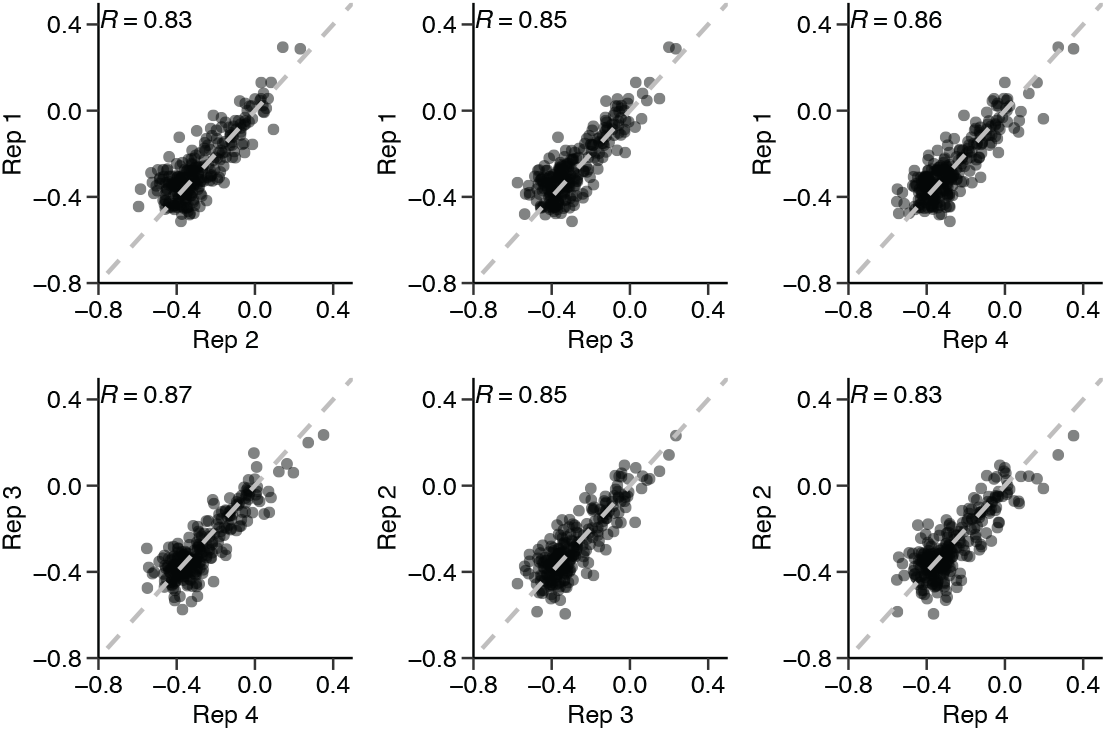
Jurkat cells exhibit strong dependence on residues in the HRD motif. Agreement of biological replicates. Fitness scores for n = 253 amino acid substitutions in the HRD-motif library are compared between biological quadruplicates. “R” is the Pearson correlation between each replicate, and the dashed line is y = x.

**Figure S5.**
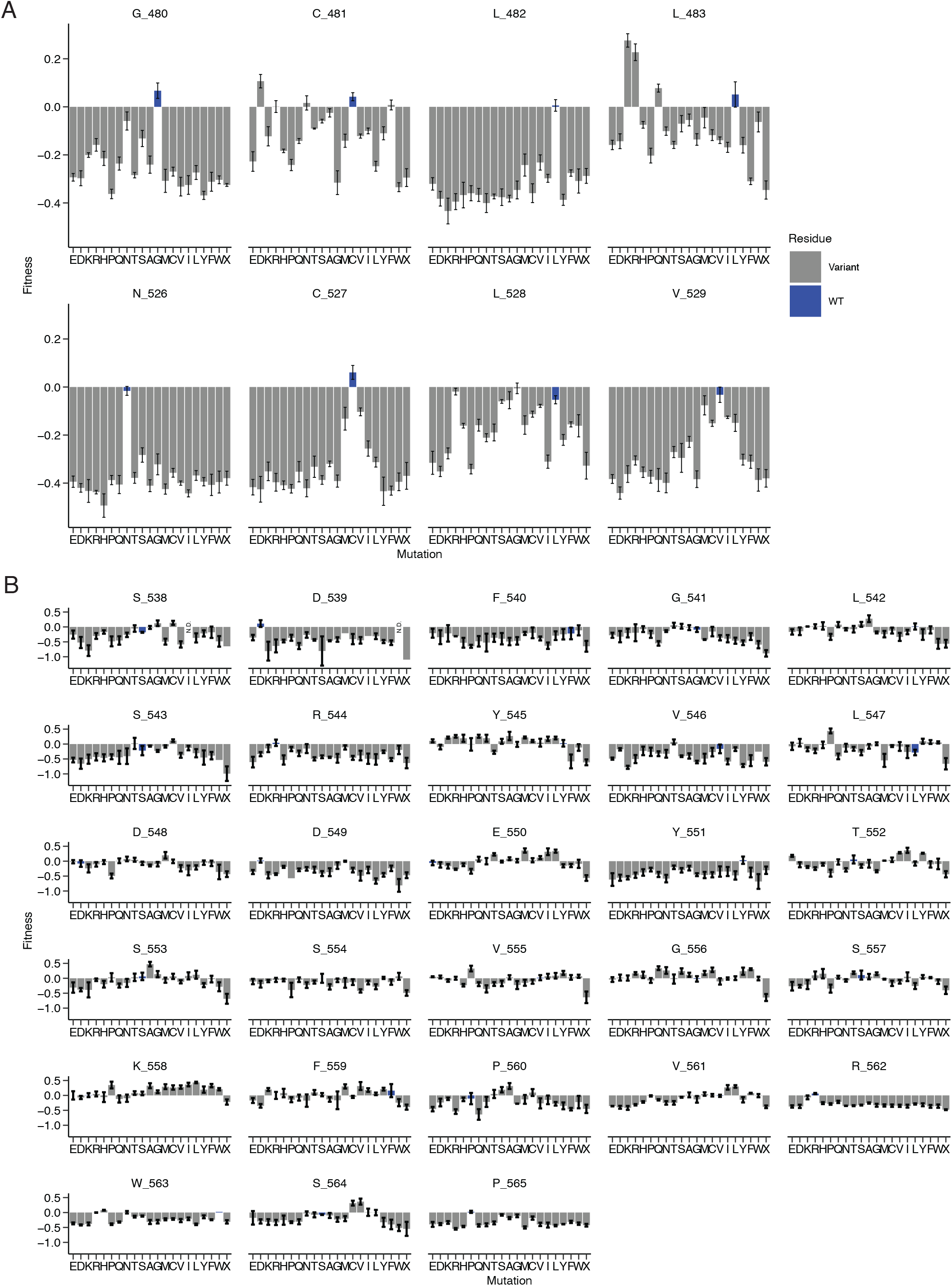
All mutations measured for the HRD and activation-loop libraries in Jurkat cells. (A) Fitness scores for all positions in the HRD-motif library in Jurkat cells. Otherwise as in Figure 2E. (B) Fitness scores for all positions in the activation-loop library. Positions where the input read counts from all four replicates failed to pass the 50-read threshold are indicated with “N.D.”. If two or three input replicates failed to pass the 50-read threshold, the mean value from the remaining one or two replicates is shown without an error bar. Otherwise as in (A).

**Figure S6.**
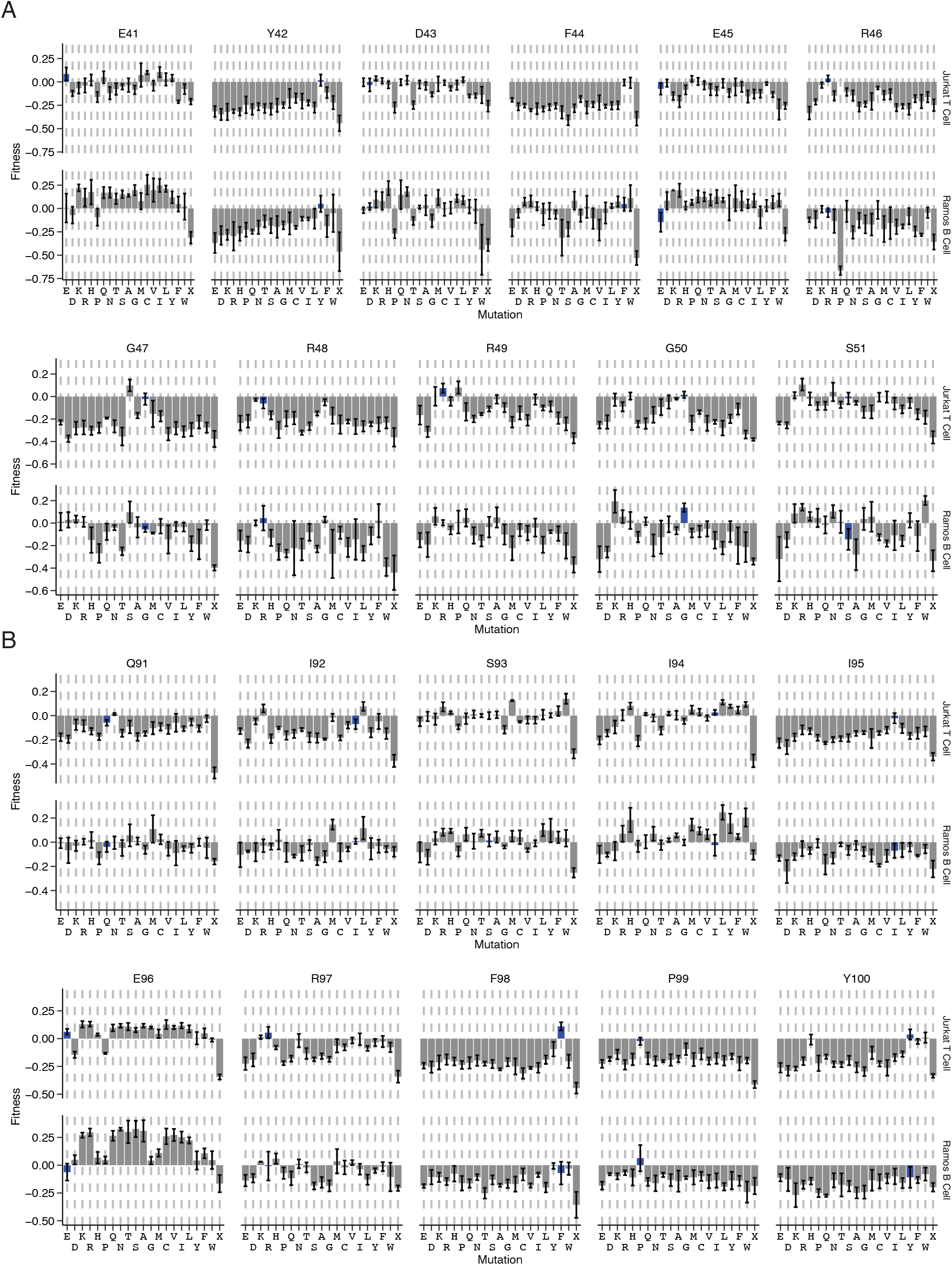
Comparison of the S1 and S2 loop libraries between Ramos cells and Jurkat cells. Fitness scores for all positions and mutations in the S1-loop library (A) or S2-loop library (B), showing the data from Jurkat cells (top row) and Ramos cells (bottom row). Otherwise as in Figure 2E.

**Figure S7.**
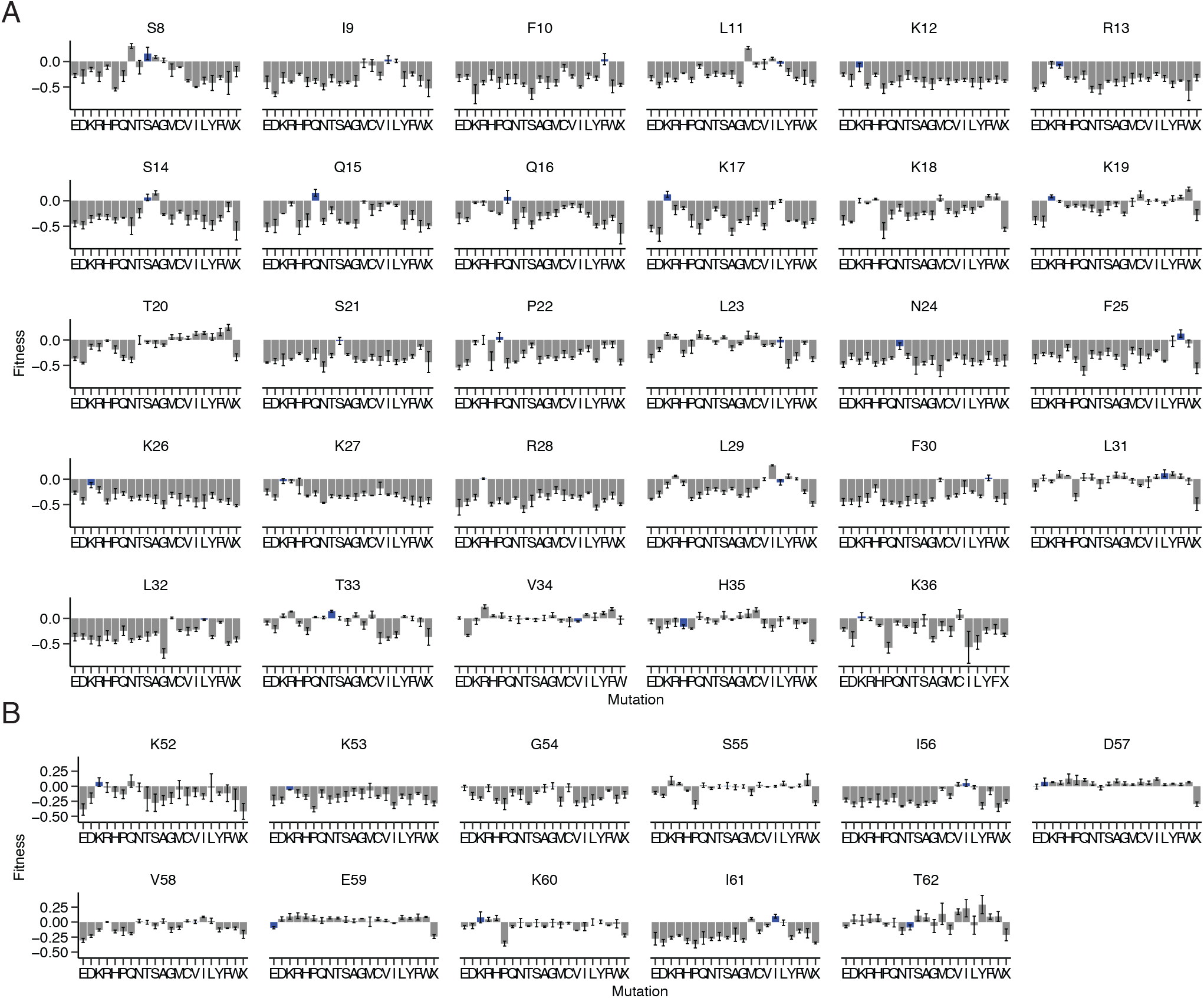
All mutations investigated for the canonical-site and peripheral-site libraries in Jurkat cells. Fitness scores for all positions in the canonical-site library (A) or peripheral-site library (B) in Jurkat cells. Otherwise as in Figure 2E. Position 37, originally designed as part of the canonical-site library, is excluded from this analysis entirely because of low read coverage for most of the substitutions.

**Figure S8.**
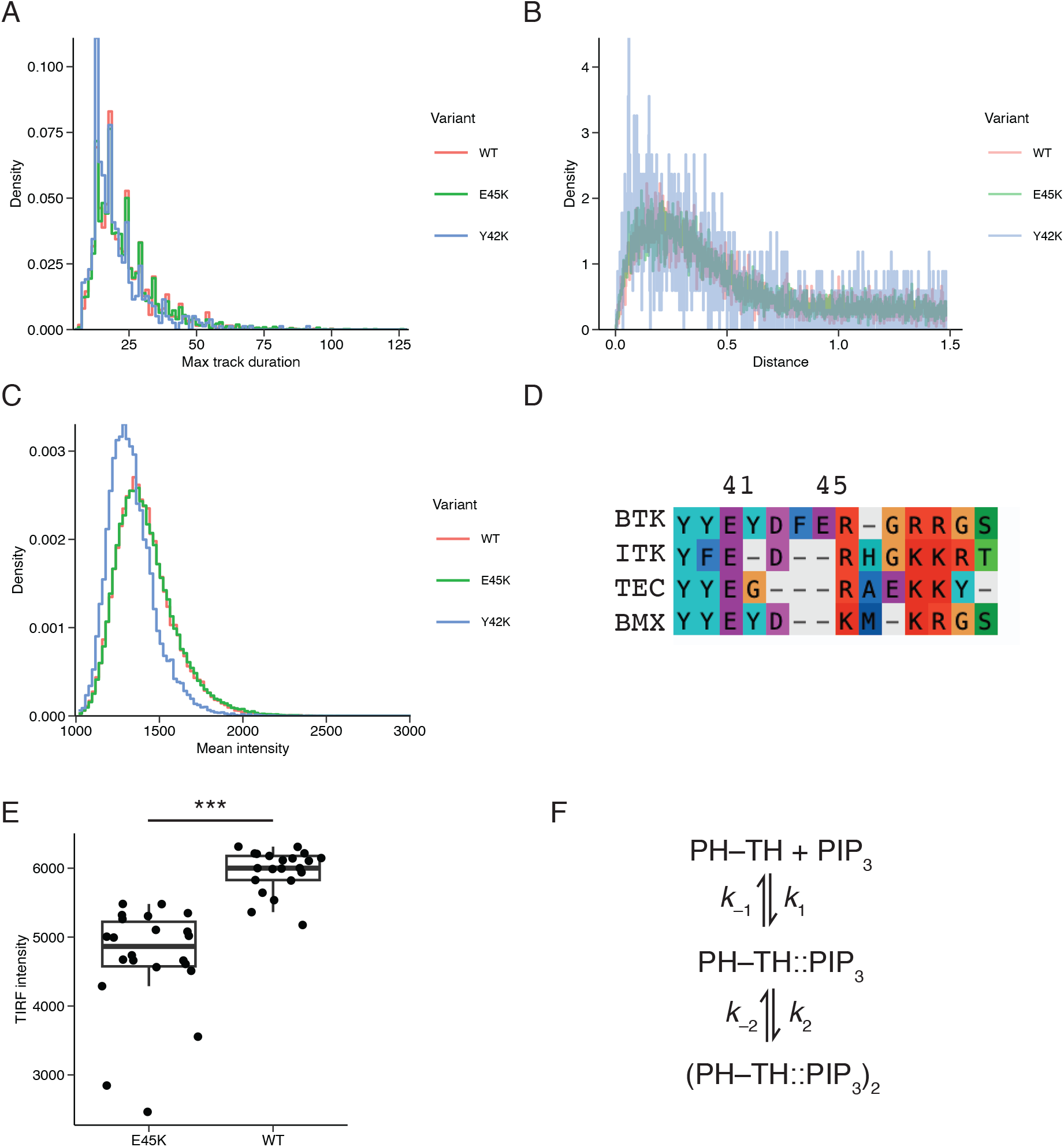
Reconstituting PH–TH domain adsorption on supported-lipid bilayers. (A) PH–TH^WT^, PH–TH^E45K^, and PH–TH^Y42K^ show the same rates of photobleaching. A histogram of the duration of the single-molecule tracks is shown for each variant (key), determined by trackmate in Fiji, on bilayers containing 4% PIP_3_. (B) Diffusion rates are the same for the three variants. The distribution of distances between 20 ms imaging steps in tracks is shown for the three variants (key). PH–TH^Y42K^ shows a noisier distribution because it has fewer tracks, as it adsorbs less to the bilayer. The spots that did not move between adjacent frames (corresponding to a distance of 0) were omitted from this and other analyses. (C) PH–TH^E45K^ and PH–TH^WT^ have the same brightness per spot. A histogram of TIRF intensity for each spot in each track is shown for the three variants (key). (D) A sequence alignment of residues 39-51 in human BTK, aligned to ITK, TEC, and BMX. Alignment was performed using MUSCLE alignment software and manually edited. The figure was generated with AliView alignment viewer. Positions 41 and 45 are indicated. (E) PH–TH^E45K^ adsorbs less than PH–TH^WT^ to bilayers containing 4% PIP_3_ at 250 pM solution concentration. PH–TH^E45K^ and PH–TH^WT^ proteins were incubated for 10 min on supported-lipid bilayers containing 4% PIP_3_ and then the median TIRF intensity across the imaging field was measured. Dots represent independent imaging fields, with the line (median), box (first and third quartiles) and whiskers (1.5x the interquartile range), with p*** < 0.001, t test. (F) The reaction scheme for kinetic modeling.

